# *In silico* transcriptomics identifies FDA-approved drugs and biological pathways for protection against cisplatin-induced hearing loss

**DOI:** 10.1101/2022.01.26.477836

**Authors:** Pezhman Salehi, Marisa Zallocchi, Sarath Vijayakumar, Madeleine Urbanek, Kimberlee P. Giffen, Yuju Li, Santanu Hati, Jian Zuo

## Abstract

Acquired hearing loss is a major health problem that affects 5-10% of the world population. However, there are no FDA-approved drugs for the treatment or prevention of hearing loss. Employing the Connectivity Map (CMap) that contains >54,000 compounds, we performed an unbiased *in silico* screen using the transcriptomic profiles of cisplatin-resistant and -sensitive cancer cell lines. Pathway enrichment analysis identified gene-drug targets for which 30 candidate drugs were selected with potential to confer protection against cisplatin-induced ototoxicity. In parallel, transcriptomic analysis of a cisplatin-treated cochlear-derived cell line identified common enriched pathway targets. We subsequently tested these top 30 candidate compounds, 15 (50%) of which are FDA-approved for other indications, and 26 (87%) of which were validated for their protective effects in either a cochlear-derived cell line or zebrafish lateral line neuromasts, thus confirming our *in silico* transcriptomic approach. Among these top compounds, niclosamide, a salicyanilide drug approved by the FDA for treating tapeworm infections for decades, protected from cisplatin- and noise-induced hearing loss in mice. Finally, niclosamide and ezetimibe (an Nrf2 agonist) exerted synergistic protection against cisplatin-ototoxicity in zebrafish, validating the Nrf2 pathway as part of niclosamide’s mechanism of action. Taken together, employing the CMap, we identified multiple pathways and drugs against cisplatin ototoxicity and confirmed that niclosamide can effectively be repurposed as an otoprotectant for future clinical trials against cisplatin- and noise-induced hearing loss.

**Significant Statement:** Employing the Connectivity Map as our *in silico* transcriptomic screening strategy we identified FDA-approved drugs and biological pathways for protection against cisplatin-induced hearing loss.

## INTRODUCTION

Platinum-based chemotherapy is a standard of care for various types of cancers, including ovarian, lung, testicular, and head and neck carcinoma (1, 2). Cisplatin, one of the most effective platinum compounds, causes permanent hearing loss in 40-60% of treated cancer patients (3–8). To reduce cisplatin damage to the inner ear cochlear cells, various therapeutic strategies including usage of antioxidants, anti-inflammatory agents, calcium channel blockers, kinase inhibitors, heat shock proteins, and thiol compounds as chemical deactivators have been used in previous studies (3–8). For example, sodium thiosulfate (STS), is effective in protecting hearing in pediatric patients with localized hepatoblastoma who received cisplatin chemotherapy; however, STS acts as a cisplatin chelator and is ineffective in protecting against cisplatin-induced hearing loss (CIHL) in patients with other types of cancers (8). Additionally, recent studies have utilized large-scale drug screens in cochlear-derived cell lines, zebrafish lateral line, or mouse cochlear explants to identify novel otoprotective compounds such as kenpaullone, and ORC-13661 (5, 8–10). To date, however, no drugs have been approved by the Food and Drug Administration (FDA) for protection against acquired hearing loss.

A promising research strategy to identify novel otoprotectant compounds is to learn from cisplatin-resistant cancer cells and try to induce the same defense mechanisms in cochlear cells. Taking advantage of a recently developed Connectivity Map (CMap) including the L1000CDS (LINCS) and Genomics and Drugs Integrated Analysis (GDA) databases (11–14), we performed *in silico* screens by connecting mechanisms of cellular resistance with therapeutic compounds associated with those biological mechanisms. The CMap consists of transcriptomic profiles of a variety of cell lines, of which many have been treated with pharmacological agents or genetic manipulations (e.g., CRISPR genomic editing). We reasoned that transcriptomic profiles favoring cisplatin resistance in the cancer cell lines should link to many drugs in the broad chemical space that are likely to induce transcriptional profiles that mimic the cisplatin-resistant phenotype, thus identifying drugs that may have novel therapeutic use for the treatment of cisplatin toxicity within the cochlea, as long as these repurposed drugs do not interfere with cisplatin’s cancer killing ability. In addition to drug identification, these transcriptomic *in silico* screens explore diverse biological pathways associated with cisplatin resistance in an unbiased manner. There are several successful studies using the CMap, including a recent study in the hearing field focusing on heat shock protein activators to treat aminoglycoside ototoxicity (9) and others repurposing existing drugs for SARS-CoV2 treatment (9, 15, 16).

In this study, we utilized transcriptomic profiles of cisplatin-resistant cancer cell lines to perform *in silico* screens, including CMap and gene set enrichment analysis (GSEA), to discover drug- and pathway-gene targets and identify compounds with otoprotective potential. Bulk RNA-seq analysis of a cisplatin-treated cochlear-derived cell line (HEI-OC1) and GSEA identified multiple common biological pathways involved in CIHL. Testing of the top 30 candidate compounds showed protective effects in HEI-OC1 cells *in vitro* and zebrafish lateral line neuromasts *in vivo* against CIHL, confirming our *in silico* screen. Niclosamide, as a top candidate and FDA-approved drug for intestinal worm infections for decades, exhibits protection against cisplatin-induced cell death *in vitro* and hair cell (HC) death in both zebrafish and mouse experimental models *in vivo*. Furthermore, we demonstrated that niclosamide also had protective effects against noise-induced hearing loss (NIHL), likely targeting common molecular pathways in CIHL and not interfering with cisplatin antineoplastic ability. Finally, we observed a synergistic otoprotective effect when niclosamide was used in combination with ezetimibe, an FDA-approved drug for the treatment of hypercholesterolemia, suggesting the possibility of a more effective multi-drug treatment for the prevention of hearing loss. Additionally, HPLC analysis, treatment of cancer cell lines *in vitro* with cisplatin and niclosamide, and previous studies in xenograft mouse tumor models have demonstrated that niclosamide does not interfere with cisplatin’s efficacy as a chemotherapeutic agent (18). Taken together, our study highlights the general use of transcriptomic *in silico* screens to identify novel therapeutics and biological pathways.

## RESULTS

Given that HCs in the inner ear are highly sensitive to cisplatin toxicity (1–8), our *in silico* approach aimed to identify small molecules capable of inducing a transcriptomic profile that could confer resistance to CIHL. For this purpose, we used publicly available RNA-seq datasets from the GEO and identified nine RNA-seq studies investigating cisplatin resistance in several cancer cell lines. In each of these studies, the transcriptomic profiles of sensitive parental cell lines were compared to those of resistant counterparts (**Figure 1-table supplement 1**). These nine studies were analyzed using the NCBI’s GEO2R tool (https://www.ncbi.nlm.nih.gov/geo/geo2r/) from which we obtained differentially expressed gene (DEG) lists. To identify compounds that will mimic the cisplatin-resistant transcriptomic profiles, we subsequently uploaded the DEGs from each study into the LINCS and GDA databases. Combined, these two CMap databases contain more than 50,000 compounds with their corresponding transcript perturbation profiles from various cancer cell lines. Our CMap database search identified more than 500 unique small molecules associated with the cisplatin-resistant phenotype. **Figure 1** (blue-shaded box) summarizes our transcriptomic-based *in silico* approach.

**Figure 1.**
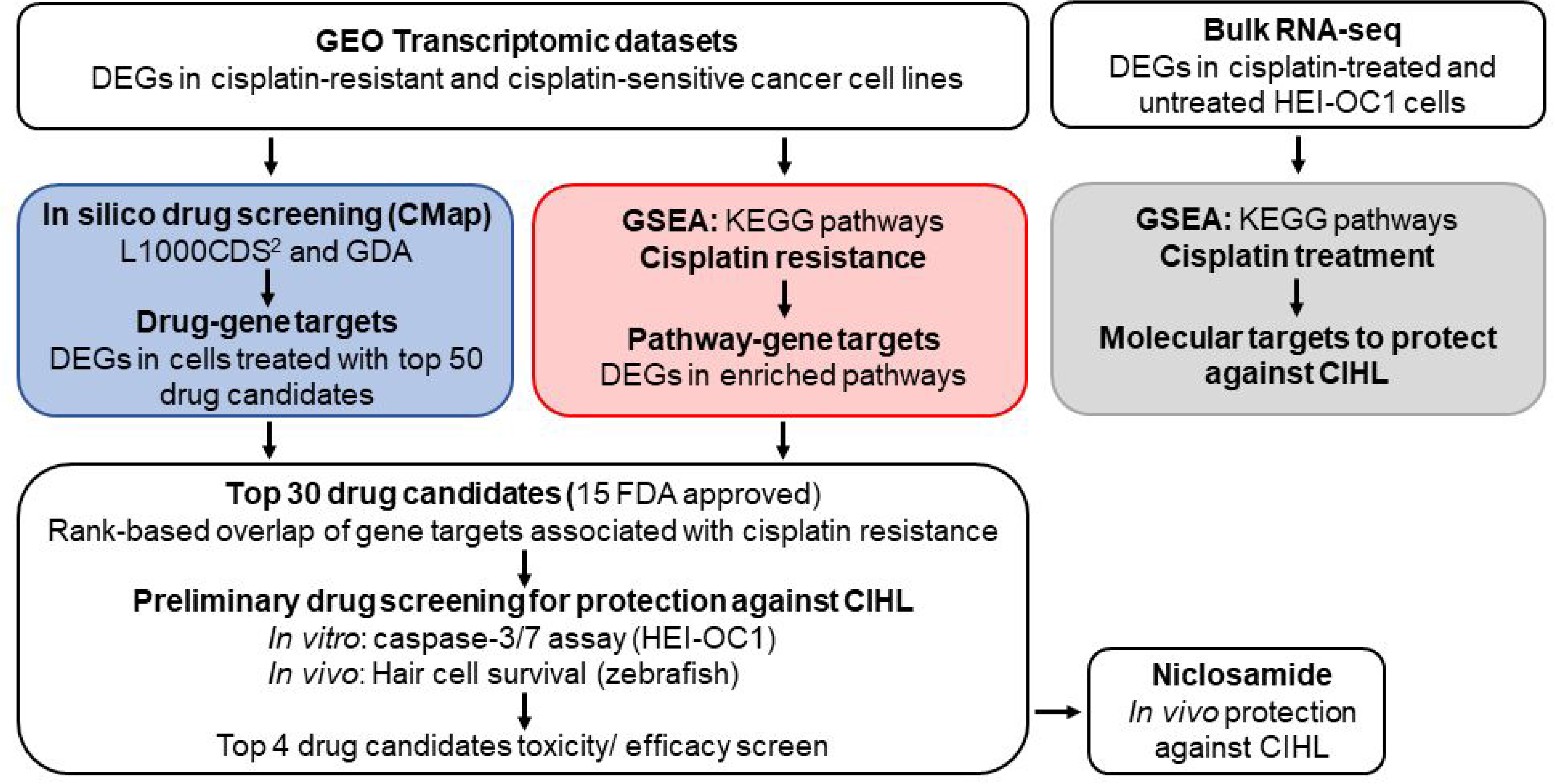
Workflow of *in silico* drug screening and signaling pathway discovery. The transcriptomic profiles of cisplatin-resistant cancer cell lines and their parental cisplatin-sensitive cells were accessed in GEO. The individual DEG lists were analyzed using the LINCS and GDA drug-gene interaction databases to identify drug-candidates that could induce the cisplatin-resistant transcriptional profile (blue-shaded box). The combined lists of up- and down-regulated DEGs were analyzed to identify enriched KEGG pathways and subsequent target genes in cisplatin resistance (red-shaded box). Drug-gene targets and pathway-gene targets were compared and used to rank the drug candidates, the top 30 drugs were validated both *in vitro* and *in vivo*, with niclosamide emerging as one of the top-hit compounds. As a complimentary approach, bulk RNA-seq of cisplatin-treated HEI-OC1 cells and GSEA analysis were performed to identify molecular targets to prevent CIHL (gray-shaded box).

In parallel to our drug screening approach, we also aimed to identify enriched signaling pathways associated with cisplatin resistance in the nine RNA-seq datasets. Each DEG list was uploaded into ShinyGO v0.66 for GSEA (17). From a total of 4,559 upregulated and 5,141 downregulated genes, the analysis identified 30 upregulated and 14 downregulated enriched signaling pathways annotated in the KEGG database (**Figure 2A** and **Figure 2-figure supplement 1**). Among the up-regulated pathways, we found the toll-like receptor, TNF, T-cell receptor, JAK-STAT, IL-17, ErbB, and chemokine signaling pathways. Additionally, the significantly down-regulated pathways included mTOR and protein processing pathways. The up- and down-regulated genes (653 and 354 genes respectively) annotated in the enriched pathways identified from GSEA were identified as pathway-gene targets for further analysis. **Figure 1** (red-shaded box) summarizes our pathway analysis approach.

**Figure 2.**
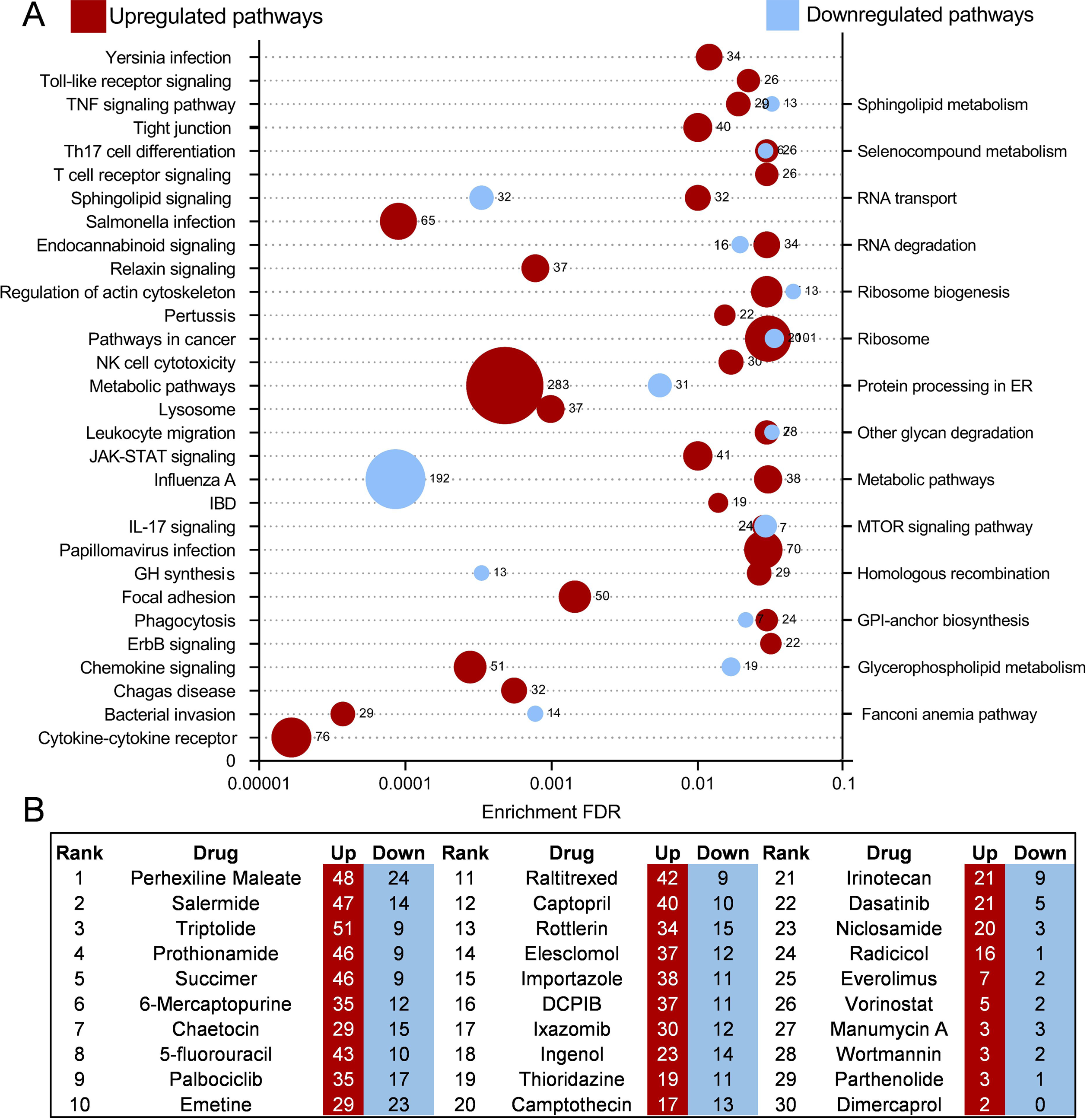
Transcriptome analysis of cisplatin-resistant cancer cell lines reveals implicated pathways and shared gene targets with identified drugs. A) Pooled cancer cell line profiles available from the GEO database were analyzed using GSEA to identify enriched molecular pathways from the KEGG database. Upregulated pathways are shown in red on the left, while downregulated pathways are shown in blue on the right. Circle size is directly correlated to the number of the genes mapped to its respective FDR value. B) Gene expression profiles for each drug derived from the iLINCS database were compared to those differentially expressed genes identified by GSEA. Overlapping genes in the same direction were then used to rank drugs. Total overlap counts for genes are included in the red up and blue down columns.

We then compared the drug-gene targets (from our drug screening) to the pathway-gene targets to rank the top compounds that may modulate the activity of specific biological pathways, and thus confer resistance to cisplatin toxicity. The drug-gene targets of each compound were retrieved from the integrated LINCS (iLINCS) portal. These drug-gene targets were then sorted into discrete up- and down-regulated gene sets and compared to the differentially expressed pathway-gene targets to rank their potential for cisplatin resistance. The top drug candidates had the greatest overlap with the genes of GSEA-identified pathways. The top 30 compounds were selected for further validation, among which 15 were FDA-approved drugs (**Figure 1**). All 30 compounds (i.e. perhexiline maleate, salermide, triptolide, prothionamide, etc.) were shown to hit at least one gene/pathway, with some drugs overlapping with more than 50 implicated genes and multiple pathways (**Figure 2B**). Niclosamide, for example, affected 20 upregulated and 3 downregulated genes in multiple pathways.

### Transcriptomic analysis of HEI-OC1 cells treated with cisplatin

To validate the relevance of cisplatin resistance in cancer cell line to CIHL, we examined the transcriptional changes associated with cisplatin treatment using the HEI-OC1 cell line derived from postnatal day 7 mouse cochleae, which has been widely used for otoprotection screenings (5, 18). Bulk RNA-seq analysis of HEI-OC1 cells exposed to cisplatin revealed differentially expressed genes despite a high degree of correlation among the cisplatin-treated and control cells (Pearson’s correlation coefficient > 0.91) (**Figure 2-figure supplement 2A**). We identified 2,728 and 1,638 genes that were significantly down-or up-regulated in the cisplatin-treated HEI-OC1 cells compared to the untreated controls (P<0.05, fold change >|1.0|) (**Figure 2-figure supplement 2B**). GSEA of differentially expressed genes in cisplatin-treated HEI-OC1 cells identified down-regulated KEGG pathways that were conversely up-regulated in the cisplatin-resistant cancer cell lines, such as the ErbB, Jak-STAT and Toll-like receptor signaling pathways, highlighting potential pathways to target for protection against cisplatin ototoxicity in cochlear cells (**Figure 1** (gray box), **Figure 2A**, **Figure 2-figure supplement 2C-E**, and **Figure 2-table supplement 2**). These independent analyses corroborate our *in silico* screens using transcriptomic profiles of cisplatin-resistant/-sensitive cancer cell lines and drug-induced genetic perturbations to identify drugs and pathways to protect from CIHL.

### Validation of top candidate drugs in HEI-OC1 cells *in vitro* and zebrafish lateral line neuromasts *in vivo*

To provide direct experimental evidence for our transcriptomic *in silico* screening, we used HEI-OC1 cells to validate our top 30 identified candidate drugs in an assay similar to our previous drug screening (5). Caspase activity was measured using Caspase-Glo 3/7 assay, as a reverse indicator of cell survival/viability. The dose responses of caspase activity for each of the 30 drug candidates were measured at various concentrations ranging from 0.002 µM to 40 µM (**Figure 3-figure supplement 1**). **Figure 3A** depicts the lowest percentage of caspase activity compared to cisplatin alone treated cells. Of the 30 compounds identified in our *in silico* screening, 20 compounds significantly reduced caspase 3/7 activity compared to controls.

**Figure 3.**
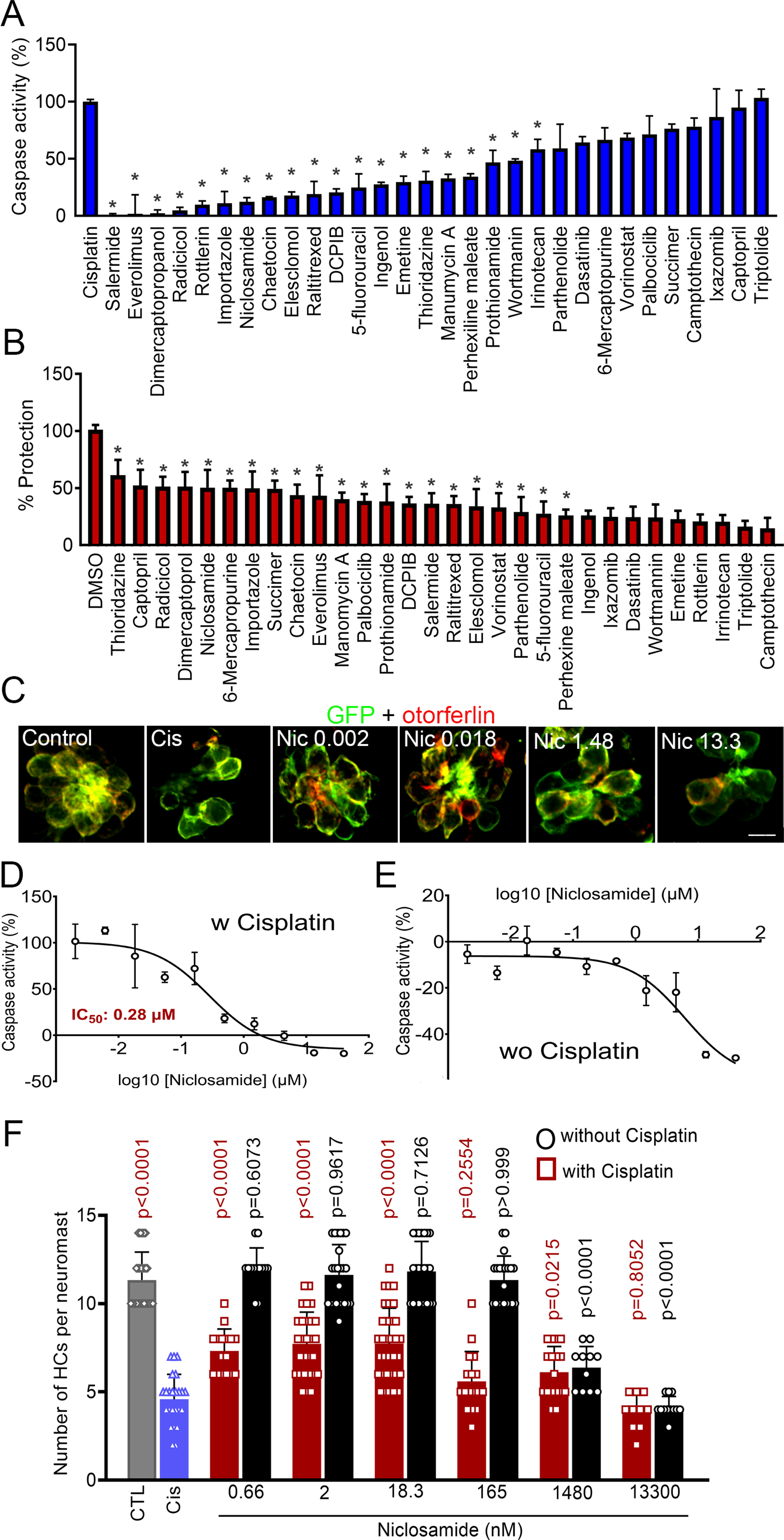
Validation of 30 experimental compounds reveals niclosamide as the top hit. A) Lowest level of caspase-3/7 activity in HEI-OC1 cells treated with cisplatin (50 µM) and experimental compounds. Caspase reads were normalized to cells treated with cisplatin/DMSO (as 100%) and cells treated with 1% DMSO (as 0%). Niclosamide reduced caspase activity to comparable levels as control cells at a dose of 4.4 µM. Data are shown as mean ± SD (n=3 wells per treatment). *P <0.05, one-way ANOVA versus cisplatin, followed by Dunnett’s multiple comparison test. B) Highest level of protection in zebrafish treated with cisplatin and experimental compounds quantified by HC count. Quantification of the HCs revealed significantly reduced cisplatin damage in zebrafish pretreated with 0.002 µM niclosamide (n=5 per group, *P <0.05, one-way ANOVA versus cisplatin-only, followed by Dunnett’s multiple comparison test). C) Representative images of zebrafish neuromasts. Niclosamide reduced HC loss at concentrations ranging from 0.002 µM-1.48 µM. GFP = green, otoferlin = red (scale bar = 20 µm). D, E) Dose response curves of niclosamide with (D) and without (E) cisplatin in HEI-OC1 cells. Results are presented as mean +/- SD. F) Niclosamide protects against cisplatin ototoxicity across multiple doses in zebrafish (n=5 per group, one-way ANOVA versus cisplatin-only (red) or versus control (black), followed by Dunnett’s multiple comparison test). Data are shown as mean ± SD.

We simultaneously tested the 30 compounds in a zebrafish *in vivo* model for cisplatin ototoxicity (19). Compound concentrations ranged from 0.002 µM to 13.3 µM. HC counts were compared to those of zebrafish treated exclusively with 300 µM cisplatin (as 0% protection) or E3 water in 0.1% DMSO (as 100% protection) to obtain the percentage protection for each compound (**Figure 3B-3C**). The drug candidates were ranked based on the most effective protection against cisplatin damage (**Figure 3B** and **Figure 3-supplement figure 2**). Of the 30 compounds, 21 showed significant levels of protection compared to zebrafish treated with cisplatin alone. When comparing the compounds showing protection in our zebrafish model (**Fig. 3B**) with the ones tested in HEI-OC1 cells (**Fig. 3A**), 15 were common in both assays and 26 (87%) in either assay. Moreover, 7 of these common compounds were FDA approved for other indications (**Figure 3-figure supplement 3**).

These experimental results strongly validated that cisplatin resistance drug-gene-pathways identified in cancer cell lines are also conserved in CIHL and therefore demonstrated the general utility of transcriptomic *in silico* drug screens.

### Niclosamide attenuates cisplatin-induced hearing loss in FVB/NJ mice *in vivo*

After our initial screening, niclosamide emerged as a potential top hit candidate based on several factors: *1)* HEI-OC1 cells treated with niclosamide reached 0% caspase activity (full protection) at a dose of ∼4.4 µM (**Figure 3A**); *2*) niclosamide provided one of the highest levels of protection (∼50%) at the lowest concentration tested (0.002 μM) in zebrafish neuromasts (**Figures 3B-C, F**); *3)* niclosamide had a relatively low calculated IC_50_ of 0.28 µM compared to other tested compounds tested in HEI-OC1 cells (**Figure 3D** and **Figure 3-figure supplement 1**); *4)* in comparison to several other hits (including thioridazine, salermide, and dimercaprol, **Figure 3-figure supplement 1**), niclosamide had a wider therapeutic window, demonstrating considerable levels of protection at over 80% of the tested doses (**Figure 3D** and **Figure 3-figure supplement 1**); *5)* niclosamide was not cytotoxic within a wide range of concentrations (**Figure 3E**); *6)* niclosamide showed levels of protection comparable to kenpaullone but better than four other benchmark compounds including sodium thiosulfate, ebselen, dexamethasone, and N-acetylcysteine (**Figure 3-figure supplement 4**) (5), and 7) niclosamide is a FDA-approved drug for the treatment of tapeworm infections for four decades with multiple clinical trials for cancer and COVID indications (NCT04753619, NCT02687009, NCT03123978, NCT04753619).

**Figure 4.**
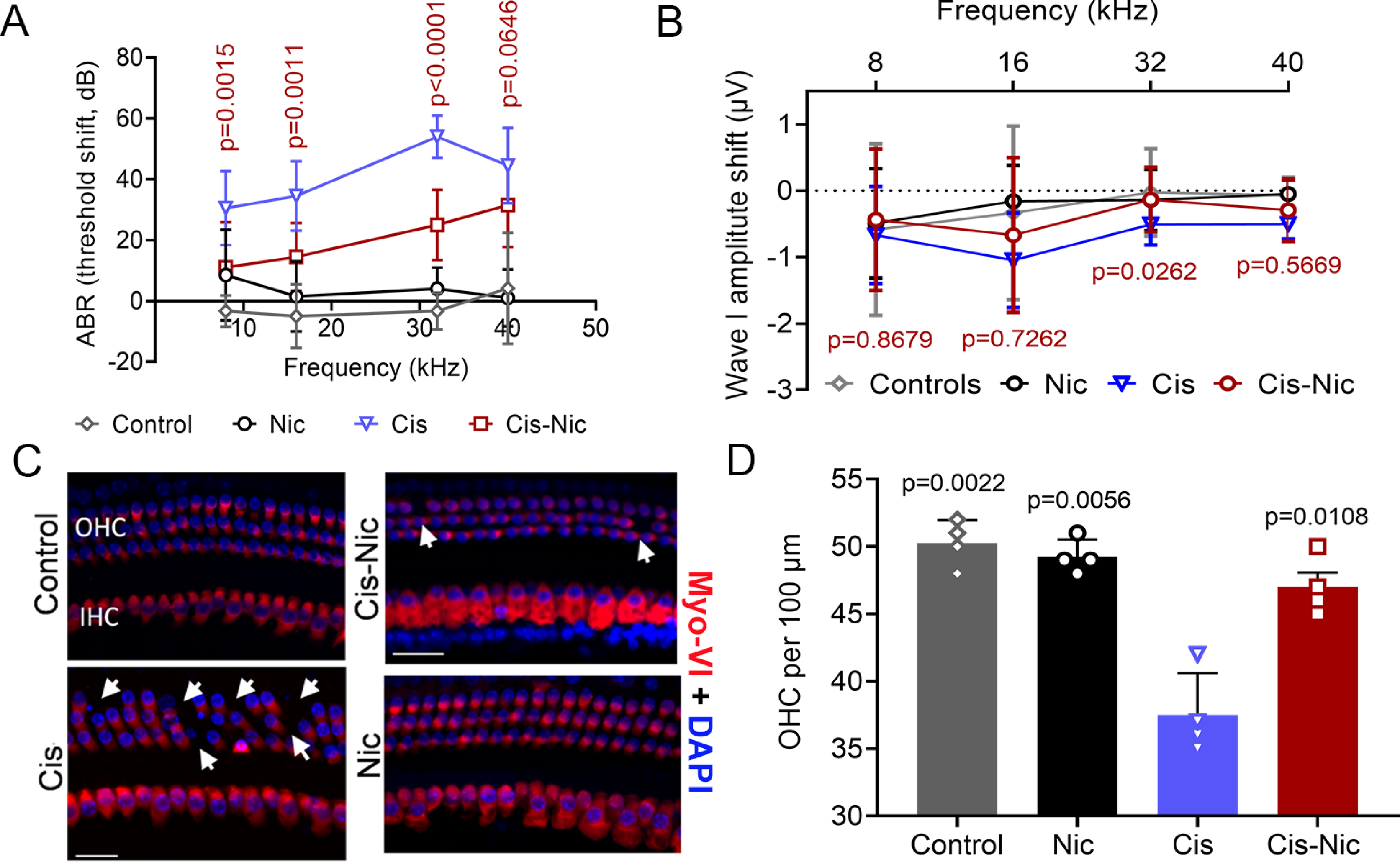
Niclosamide demonstrates otoprotective effects against cisplatin *in vivo*. A) Niclosamide-treated animals have significantly reduced ABR threshold shifts at 8, 12, and 32 kHz as compared to cisplatin-only treated mice (n=8 per group, two-way ANOVA versus cisplatin-only treatment, followed by Tukey’s multiple comparison test). B) Wave I amplitude shifts. Animals exposed to cisplatin and treated with niclosamide showed a significant reduction in wave I amplitude shifts at 32 kHz compared to cisplatin-only treated animals (n=8 per group, one-way ANOVA versus cisplatin treatment, followed by Tukey’s multiple comparison test). C) Representative immunofluorescent images of the mid-basal region of the cochlea stained for Myosin-VI (red) and DAPI (blue) showing minimal levels of hair cell loss when animals were cotreated with cisplatin and niclosamide compared to cisplatin-only treated animals. White arrows denote missing hair cells (scale bar = 20 µm). D) Quantification of outer hair cells from immunofluorescent images shows that cotreatment with niclosamide grants full protection against cisplatin-induced hair cell loss (n=5 per group, ,one-way ANOVA versus cisplatin, followed by Dunnett’s multiple comparisons test). Data shown as mean ± SD.

Thus, we decided to move forward with the characterization of niclosamide’s protective effect in a mouse model for CIHL. For this purpose, mice were randomly assigned to four different treatments: control (vehicle), cisplatin-alone (cisplatin + vehicle), cisplatin + niclosamide, and niclosamide-alone. Niclosamide (10 mg/kg for 4 consecutive days) and cisplatin (30 mg/kg single day divided into two doses) were injected IP. The results of the ABR tests at day-5 post cisplatin injection showed a statistically significant difference between the hearing threshold shifts of cisplatin-niclosamide treated mice at 8, 16, and 32 kHz (P < 0.05) when compared to cisplatin only group (**Figure 4A** and **Figure 4-figure supplement 1**). The ABR thresholds of niclosamide-only treated mice were not significantly different from saline-injected controls. These *in vivo* ABR results show that niclosamide protects against CIHL in mice.

The ABR wave-1 amplitude represents the summed activity of the cochlear nerve, and therefore, an informative measure of auditory synapse function. We measured mean wave-1 amplitudes at 8, 16, 32, and 40 kHz, in the control and niclosamide-treated mice before cisplatin injection and at day-5 post-cisplatin injection. Wave I amplitude shifts from the 85 dB SPL stimuli were compared between groups using a two-factor ANOVA (group x frequency). At day 5 post-treatment, the two-factor ANOVA revealed a significant two-way interaction of group x stimulus level at 32 kHz and the post-hoc Tukey’s test revealed that the cisplatin-niclosamide treated group had a reduced wave I amplitude shift compared to cisplatin-only group (**Figures 4B** and **Figure 4-figure supplement 2**). These ABR wave-1 amplitude results provide further evidence of niclosamide’s otoprotection *in vivo*.

We further quantified the number of outer HCs (OHCs) at the mid-basal region, the most protected frequency region shown by ABR threshold and wave I amplitude measurements. Representative images of cochlear HCs are displayed in **Figure 4C**. Quantitative data for HC count at the mid-basal region are displayed in **Figure 4D**. The one-way ANOVA revealed a significant group effect (P <0.05). The post-hoc test revealed that while inner HC survival was not affected, the cisplatin-niclosamide group had more OHC survival than the cisplatin alone group in the mid-basal region (**Figure 4D**). These ABR and HC count data together, confirmed that niclosamide protects OHCs against cisplatin damage.

### Niclosamide protects NMDA-induced HC loss in zebrafish *in vivo*

Since CIHL and NIHL share mechanistic commonalities (23, 24), we examined whether niclosamide had any protective effects in a zebrafish model for HC excitotoxicity (25). As previously described, neuromast HC numbers were reduced after exposure to 300 μM NMDA (22). Conversely, post-treatment of the zebrafish exposed to 300 µM NMDA with 0.002 µM or 0.0183 µM of niclosamide resulted in significantly increased HC survival (**Figure 5A**). These zebrafish excitotoxicity results indicate that niclosamide may also protect against NIHL.

**Figure 5.**
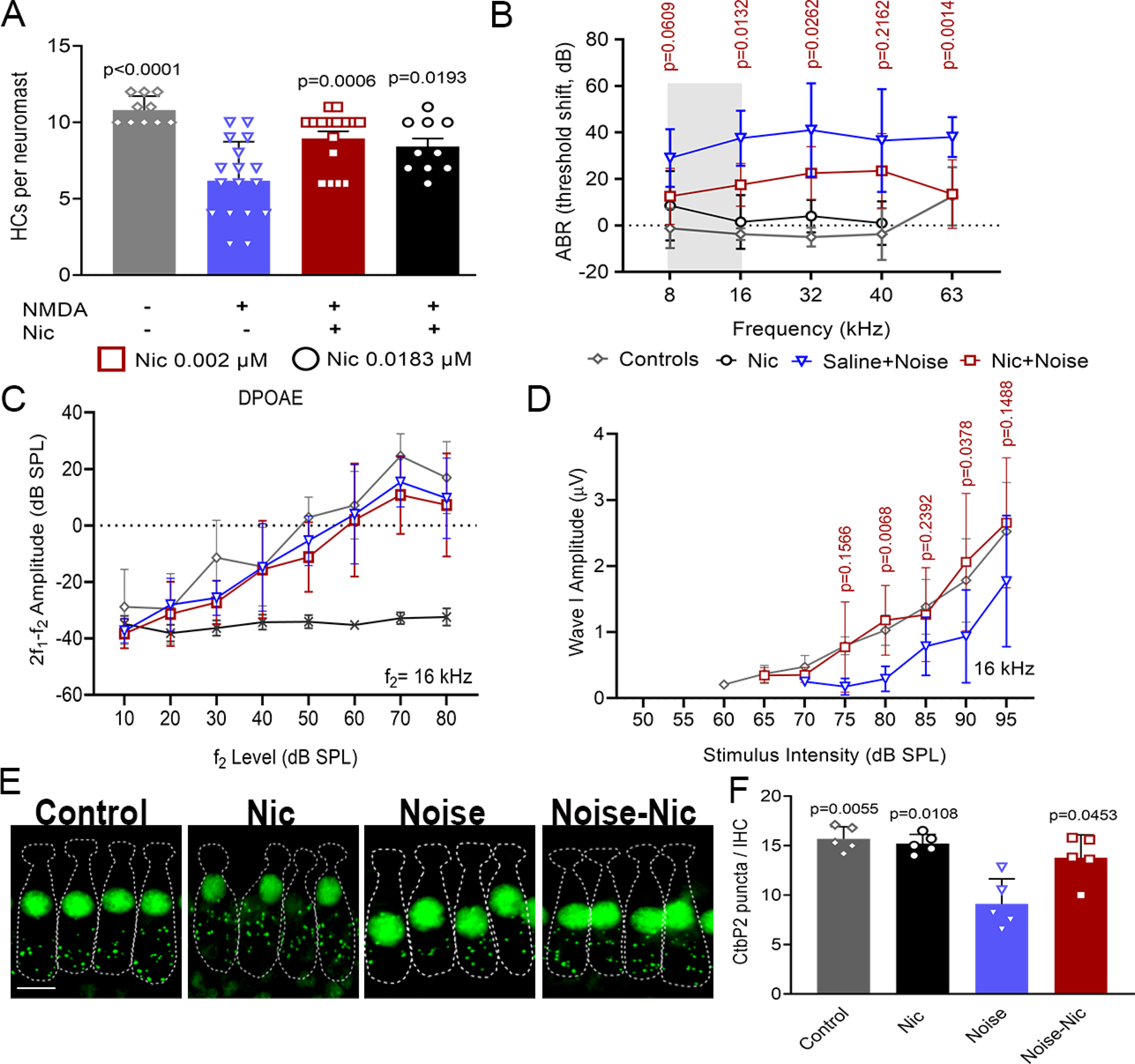
Niclosamide protects against NIHL. A) Niclosamide reduces NMDA excitotoxicity in zebrafish neuromasts. Zebrafish were treated with 300 µM NMDA (25) followed by 0.002 or 0.0183 µM niclosamide. At both doses tested, niclosamide’s treatment showed a significantly increase in the number of hair cells compared to NMDA-only treated zebrafish. (n=5 per group, p values were calculated against NMDA-only, one-way ANOVA, Dunnett’s multiple comparisons test). B) Noise-exposed mice treated with niclosamide had significantly lower ABR threshold shifts compared to saline-treated animals (n=8 per group, two-way ANOVA versus noise-only treatment, followed by Sidak’s multiple comparisons test). C) There were no differences in DPOAE amplitudes across all groups from 10-70 dB SPL (n=8 per group, two-way ANOVA, followed by followed by Sidak’s multiple comparisons test). D) Niclosamide-treated mice showed comparable wave-1 amplitudes across 65-90 dB SPL to age-matched controls and significantly higher wave-I amplitudes at 80- and 90-dB SPL than saline and noise-exposed mice (n=8 per group, two-way ANOVA, followed by Tukey’s multiple comparisons test versus noise). E) CtBP2 staining (green) showed that niclosamide protects against synaptic loss after noise exposure. F) Quantification of CtBP2 puncta per inner hair cell (n=4 per group, one-way ANOVA versus noise-only treatment, followed by Dunnett’s multiple comparisons test). Data shown as mean ± SD.

### Niclosamide diminishes NIHL in adult FVB/NJ mice *in vivo*

We further investigated niclosamide’s therapeutic effects against NIHL in FVB/NJ mice. We first injected the mice with 10 mg/kg niclosamide via IP once per day for four consecutive days, starting one day before noise exposure, the day of the noise exposure, and two more days after noise exposure. Control animals received vehicle injections on the same schedule. Noise exposure was administered at 8-16 kHz at 100 dB SPL for 2 hrs. Noise-induced ABR threshold shifts were obtained by subtraction of the pre-exposure from the post-exposure thresholds. Two-way ANOVA followed by Sidak’s multiple comparison test revealed that the niclosamide-noise exposed group had lower threshold shifts than noise-exposed group across most of the tested frequencies (16 kHz, 32 kHz and 63 kHz) at day 14 (**Figure 5B** and **Figure 4-figure supplement 1**). These results demonstrate that niclosamide also protects against NIHL in mice and suggest that its action is independent of cisplatin inactivation.

To determine whether niclosamide prevents NIHL by protecting OHCs, we measured the DPOAE amplitudes at the different f_2_ frequencies with L2 levels ranging from 10 to 70 dB SPL (**Figure 5C**). In the noise-niclosamide group, DPOAE amplitudes were not significantly higher than the noise-saline group at day 15 post-noise exposure. A two-factor ANOVA (group x frequency) was used to compare pre-exposure amplitudes to day 15 amplitudes. The ANOVA revealed no significant two-way group x frequency interaction indicating that the OHC function is similar between all groups and suggesting that niclosamide’s protective effect against noise could be due to prevention of synaptopathy between inner HCs and cochlear nerves. To test our hypothesis, mean ABR wave-I amplitudes at 8, 16, 32, and 40 kHz were measured at day 15 post-noise exposure. Amplitudes from the 10 to 90 dB SPL stimulus intensity were compared between groups in the pre-noise test using a two-factor ANOVA (group x stimulus level), and no group differences were detected (data not shown). At day 15, only the 50-90 dB SPL stimulus levels were used because many of the subjects had no responses below 50 dB SPL. Results from these experiments showed that the wave I amplitudes from the niclosamide-noise group were increased at all the noise stimulus tested, with 80 and 90 dB SPL showing significant differences. The two-way ANOVA revealed a significant interaction of group x stimulus level (P<0.001). The Tukey’s post-hoc revealed that the niclosamide-noise group had higher amplitudes at 80 and 90 dB SPL compared to the noise-exposed group (**Figure 5D**). The ABR wave-I amplitude results showed that cochlear nerve activity in the noise-niclosamide group was comparable to the aged-matched controls, with no statistically significant difference between these groups, thus providing evidence of niclosamide’s protection from synaptopathy.

To assess the protection of the ribbon synapses, the cochlear samples were immunostained with CtBP2. Representative images of the mouse ribbon synapses at 16 kHz are displayed in **Figure 5E**. Quantitative data for ribbon synapses at 16 kHz are displayed in **Figure 5F**. T-test statistical analyses revealed that the niclosamide-noise group had more synaptic ribbons than the saline-noise group (**Figure 5F**). The frequency region of 16 kHz was used for CtBP2 ribbon count because it has been shown that ribbons are more abundant in this frequency region (26).

Taken together, our results showed that niclosamide protects against CIHL and NIHL in both zebrafish and mice *in vivo*. Its protection in mice is prominently represented by OHC survival in CIHL (**Figure 4C-D**) and ribbon synapse protection in NIHL (**Fig. 5E-F**). However, it is very likely that niclosamide might be also exerting its protective effect on other cochlear cells.

### Niclosamide shows synergistic effects with the Nrf2 agonist ezetimibe in zebrafish

Given the multiple pathways that niclosamide affects (20, 27-31, and **Figure 2B**) and the key role of Nrf2 in regulating reactive oxygen species (ROS) in cisplatin toxicity (**Figure 2-figure supplement 1**, 31), we reasoned that niclosamide could synergize with activators of the Nrf2 pathway and thus increase the levels of protection against cisplatin ototoxicity. For this purpose we used the zebrafish model for cisplatin ototoxicity to test niclosamide’s protective effect in the presence of ezetimibe, a potent Nrf2 activator and FDA-approved cholesterol-lowering medication (32, **Figure 6A**). The synergistic effect of the combination of niclosamide and ezetimibe at different concentrations was determined using classical synergy models (Bliss and Loewe) implemented in the program Combenefit (33, 34). Both the Bliss and Loewe models suggested highest synergistic effect between niclosamide and ezetimibe in the prevention of cisplatin damage to zebrafish HCs when used at 1.48 µM ezetimibe and 0.66 nM niclosamide (**Figure 6B**). Other dose combinations showing synergy are shown in dark blue boxes in the synergy matrix plot (**Figure 6C**). These results demonstrate that niclosamide and the Nrf2 activator, ezetimibe, act in synergy against cisplatin ototoxicity through activation of the Nrf2 pathway, while niclosamide could act through multiple Nrf2-independent pathways such as those identified in our pathway analysis of cisplatin-resistant cancer cell lines (**Figure 2**, **Figure 2-figure supplement 2 and Figure 2-table supplement 2**).

**Figure 6.**
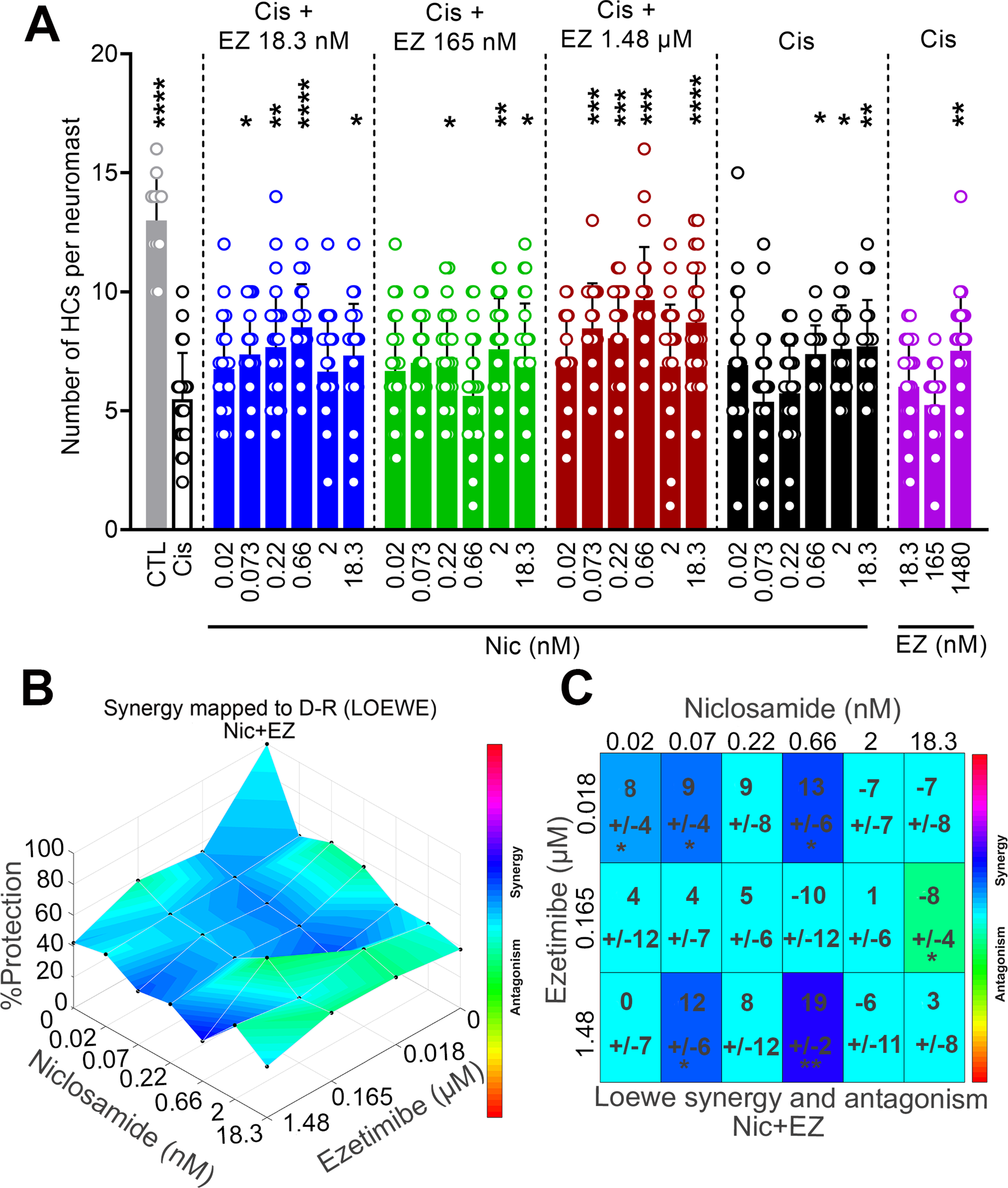
Niclosamide shows synergistic effects with the Nrf2 agonist, ezetimibe. A) Cotreatment of niclosamide with the Nrf2 agonist, ezetimibe, demonstrates an increased in hair cell protection. Zebrafish were treated with combinations of niclosamide (0.02-18.3 nM) and EZ (0.0183-13.3 µM). Ezetimibe alone + cisplatin showed higher HC counts at 1.48 µM, while niclosamide alone + cisplatin showed higher hair cell counts at concentrations equal or lower than 2 nM. However, combining both compounds showed significantly higher hair cell counts across a much lower range of doses for both niclosamide and ezetimibe (n=5 per group, p values were calculated against cisplatin-only, one-way ANOVA, followed by Dunnett’s multiple comparisons test). B, C) Niclosamide and ezetimibe show synergistic/additive otoprotection in zebrafish. A three-dimensional plot (B) showing dose response protection in zebrafish. Loewe synergy and antagonism scores (C) calculated for each combination of doses indicate a highest synergistic activity with 0.66 nM niclosamide and 1.48 µM ezetimibe (n=5 per group). Other dose combinations showing synergy are shown in dark blue boxes. Dose combinations with scores of 0 and 1 show additive effect. *P<5x10^-2^; **P<10^-3^, ***P<10^-4^ versus control fish, One-sample t-test run by the Combenefit software (33, 34). Data is shown as mean ± SD

### HPLC analysis demonstrates no interaction between niclosamide and cisplatin

Drug-drug interactions (DDI) through chemical binding could have a negative impact on cisplatin’s antineoplastic effects. A simple explanation of niclosamide’s protection against CIHL is that it can directly inactivate cisplatin, similar to several otoprotectants (e.g. sodium thiosulfate) that are currently in clinical trials (7, 8). Although our results on NIHL and additional xenograft mouse cancer model studies strongly suggest otherwise, we further investigated any possible DDI between niclosamide and cisplatin. First, by developing an HPLC method, we showed no chemical interaction between niclosamide and cisplatin (absence of third peak) at several dose ratios of niclosamide and cisplatin (**Figure 7A**). Second, by employing the seminoma cancer-derived cell line, NCCIT, in *in vitro* experiments, we demonstrated that niclosamide does not interfere with cisplatin tumor killing activity (**Figure 7B**). Overall, our *in vitro* results demonstrated there is no chemical or biological interaction between niclosamide and cisplatin, and that niclosamide is acting as a therapeutic compound to prevent not only CIHL but also NIHL. Moreover, these last results are consistent with the previously described synergistic cancer killing activity between niclosamide and cisplatin in renal cell carcinoma (RCC) xenograft models (18).

**Figure 7.**
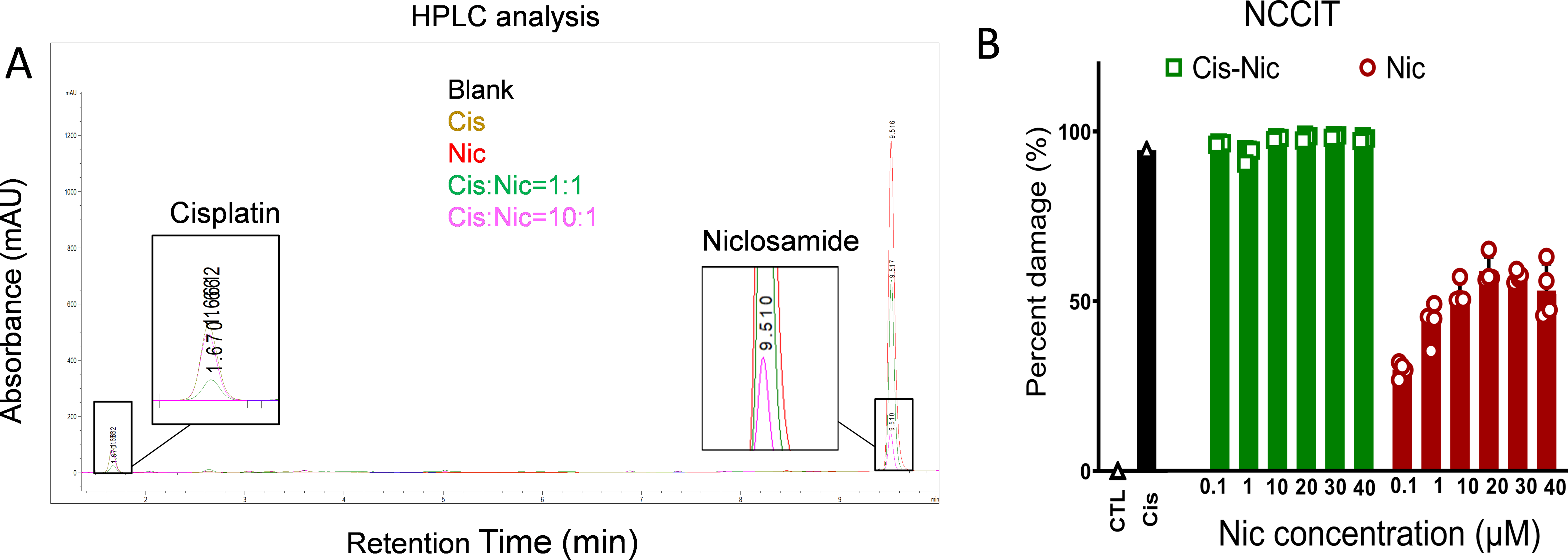
Niclosamide does not interact with cisplatin. A) HPLC analysis demonstrate that there is no chemical interaction between cisplatin and niclosamide at various concentrations. B) NCCIT cancer cells were used in a viability test to show that niclosamide (0.1-40 µM) does not inhibit cisplatin anti-cancer activity.

## DISCUSSION

Hearing loss caused by cisplatin, noise, antibiotics, and aging affects 5-10% of the world’s population (35, 36). To date, no drugs have been approved by the FDA for clinical use to prevent such types of ototoxicity. In this study, we applied widely used bioinformatics tools (CMap) to cisplatin-resistant and -sensitive cancer cell lines and performed transcriptomic *in silico* screens of over 54,000 compounds in the CMap. We identified 44 pathways and more than 30 drug candidates, most of which have never been previously associated with otoprotection. By employing the inner ear HEI-OC1 cell line and zebrafish neuromasts, we validated the top 30 compound hits, 26 of which exhibited protection in either assay, confirming our *in silico* screens. We then zeroed in on niclosamide, a previously FDA-approved drug that has been widely used in humans for tapeworm treatment for decades. In addition to its excellent pharmacokinetic/dynamic (PK/PD) properties and safety profile (27, 37–39), niclosamide exhibits outstanding protective effects against cisplatin- and noise-induced hearing loss in both zebrafish and mice when administered systemically. In summary, our work highlights that *1)* by using the CMap it is possible to identify compounds that can regulate biological pathways associated with various pathogenic conditions including acquired hearing loss; *2)* FDA-approved drugs can be repurposed to prevent cisplatin- and noise-induced hearing loss in clinics; and *3)* by targeting multiple biological pathways (37–39) rather than individual pathways, niclosamide can exert better protection than targeting individual pathways against acquired hearing loss.

### Transcriptomic *in silico* screens for therapeutic drugs and biological pathways for hearing loss

The concept of the CMap is to connect molecular pathway genes and pharmaceutical drugs through transcriptomic profiles. A large collection of transcriptomic profiles has been obtained and categorized from a variety of cell lines and tissues that have also been genetically manipulated or treated with each of the approximately 54,000 compounds included within these databases. Such databases can be unbiasedly screened for both molecular pathway genes and drugs that exhibit similar or opposing transcriptomic profiles. The CMap has been used for transcriptomic *in silico* screens for other physiological phenotypes (9, 11, 12, 15, 16). Clearly, transcriptomic *in silico* screens do not require high-throughput facilities and thus can be widely used. What are needed for the use of transcriptomic *in silico* screens, however, are specific transcriptomic profiles of the two conditions in question, either in cells or in tissues. Our approach here provides a successful example on how to use such powerful tools (CMap) to identify drug candidates for further validation in subsequent assays. More importantly, such approaches can be effectively applied to situations where no cell line assays are appropriate for drug screens. For example, drug screens for NIHL cannot be performed physically in cell lines; we envision that such *in silico* screens would be ideal due to the availability of several cochlear transcriptomic profiles for NIHL in animal models (40, 41). There are no immediate prospects of drug candidates to be approved by the FDA for protection against NIHL, antibiotic-induced or age-related hearing loss, which affect much larger populations than CIHL, the CMap can therefore be fruitful in these important unmet health arenas.

Although CMap has been a popular resource for data-driven drug repositioning using a large transcriptomic compendium (42), the genes-pathways-drugs identified using CMap demand further validation in relevant experimental assays. The extensive overlap between the molecular signatures associated with cisplatin resistance in cancer cells and cisplatin ototoxicity in HEI-OC1 cells (**Figure 2-figure supplement 2**) provides validation for our screening approach. Furthermore, many pathways we identified have been previously implicated in otoprotection (i.e., B-Raf, CDK2, STAT3 and others) (5, 43). Iterative ranking of drug candidates based on overlap of target genes in enriched molecular pathways of cisplatin resistance allowed for unbiased selection of compounds for further testing. Importantly, in two widely used CIHL assays, 26 of our top 30 drug candidates tested all exhibited protection in at least one assay. Further *in vivo* testing showed that a top candidate, niclosamide, protects against CIHL in mice.

### Repurposing FDA-approved drugs for treatment of hearing loss

Among the 30 drug candidates we identified and validated in our *in silico* screens, 15 are FDA-approved for indications other than otoprotection. Recently, focused preclinical studies testing of specific FDA-approved drugs for hearing protection (e.g., statins) have yielded satisfactory results (5, 43–45), thus making these 15 new FDA-approved drugs attractive for repurposing as otoprotectants.

In general, repurposing FDA-approved drugs offers many advantages over developing new chemical entities (NCE) (46). The first and most significant is that the safety and PK/PD profiles of FDA-approved drugs are well-defined in their respective dosing and formulation requirements in both preclinical and clinical studies. With such data available, even drugs conventionally considered to have undesirable safety profiles may be repurposed at tolerated dosages for otoprotection (46, 47). This also applies to our top drug candidate, niclosamide, which originally was approved by the FDA for its toxic effect against tapeworm infections in humans before later being shown to protect kidneys against oxaliplatin damage at lower plasma levels compared to the gastrointestinal system (27, 30). The second significant advantage of repurposing FDA-approved drugs is the fast and cost-effective path to clinics. Based on FDA data since 2003, the number of approved repurposed drugs has surpassed that of NCEs (46).

### Niclosamide as a candidate drug for clinical trials on hearing loss

With an already established safety profile in humans, niclosamide serves as a promising therapeutic candidate for expedited FDA approval. Niclosamide has been used safely for nearly 40 years since 1982 for the treatment of parasitic infections in humans (20, 27, 28). Although it has been discontinued in the US due to commercial reasons, it represents an excellent opportunity to repurpose this FDA-approved drug for new indications such as treatment for cancers and hearing loss (37, 48, 49). Niclosamide has many desirable drug properties: (a) it is effective by itself against colon metastatic tumors such as HCT116, SW620, LS174T, SW480, and DLD-1 and is synergistic with cisplatin in a xenograft mouse model of renal cell carcinoma through Wnt/β-catenin signaling (20, 30), it can be delivered orally, which is a significant advantage over other local delivery methods. Our results support that niclosamide can cross the BLB in mice. Niclosamide also has well-defined safety and PK/PD profiles in nonclinical and clinical use. Current ongoing clinical trials for cancer treatment commonly use oral doses up to 2 g/day without any adverse effects (48, 49). This dose was found to lead to serum C_max_ ranging from 0.8 to 18 μM (37). In CD1 mice, IP injection of niclosamide at 10 mg/kg single dose (the otoprotective dose for our NIHL and CIHL) resulted in a C_max_ of 40 μM (38). In contrast, 120 mg/kg single dose of niclosamide via oral gavage in mice led to a serum C_max_ at 2 μM at 1 hour, suggesting oral delivery in mice is not optimal (lower bioavailability than in humans) (39). Despite the low bioavailability of the original drug formulation that has been used over 40 years, for the purpose of cellular protection, a drug concentration lower than those approved for cancer treatment might be needed. In support, the IC_50_ values of niclosamide in our *in vitro* cell line assay (0.28 μM) and *in vivo* zebrafish assay (<0.002 μM) (**Figure 3**) are 100-9,000x fold less than 18 μM (serum C_max_ corresponding to 2 g/day human dose), supporting that niclosamide should be sufficiently potent even if it is only partially absorbed from the intestinal tract. Alternatively, new formulations of niclosamide can be further tested to increase bioavailability through oral administration or bypassing gastrointestinal tract through intramuscular or intravenous injections.

### Niclosamide and the possible mechanisms of action for protection against cisplatin ototoxicity

Although a number of pathways have been implicated in other tissues since its original discovery as an essential world-wide medicine, niclosamide’s otoprotective mechanism was never explored. Here we demonstrated that while several potential downstream pathways are likely affected in the cochlea upon niclosamide treatment, synergistic effects are observed when co-incubated with Nrf2 agonists in zebrafish; suggesting that a combinational treatment at lower doses will be more beneficial to treat hearing loss than individual niclosamide’s treatments at higher doses. Thus, targeting multiple pathways is more effective than targeting a single pathway in battling ototoxicity, and that might be the key for niclosamide’s success. Interestingly, both niclosamide and ezetimibe have been widely used for different pathological conditions, thus their combination would have an exciting potential for safe treatment of ototoxicity. Similarly, given that statins are effective in otoprotection (50) and ezetimibe has a similar indication as statins in lowering cholesterol, our results suggest that the combination of niclosamide with statins could have better otoprotection than single-drug treatment.

Though niclosamide serves as an ideal potential therapeutic for repurposement for FDA approval for both cisplatin- and noise-induced hearing loss, future studies should be used to identify the exact mechanism under which it exerts its otoprotective effects. Given that our study has elucidated a number of potentially implicated pathways, many of which have already been identified in previous studies into hearing loss, we posit that the therapeutic benefits of niclosamide may arise from a large number of mechanisms that are all converging on cell survival pathways that are typically downregulated in HCs by cisplatin treatment and noise injury. While numerous converging pathways will likely make the process of identifying an exact mechanism complicated, such a mechanism may explain niclosamide’s robust protection against two distinct insults with divergent mechanisms. While *in silico* gene set enrichment analysis served to reveal a number of these potential otoprotective pathways, each of these pathways must be validated *in vitro* and *in vivo* to determine exactly what role they may have in granting niclosamide’s otoprotection. Though the exact molecular mechanism remains unknown, the identification of niclosamide as a novel otoprotective agent nevertheless demonstrates the advantages of using the connectivity map to conduct large-scale drug screens, the results of which can be further reinforced by both *in vitro* and *in vivo* studies to identify and characterize novel drug candidates.

## MATERIALS AND METHODS

### Materials

All drugs tested were purchased from Cayman Chemical (USA). Cisplatin vials (aqueous solution of 1 mg/mL, Accord Healthcare, Durham, NC) were obtained from Creighton University Pharmacy.

Antibodies used included: C-terminal binding protein-2 (mouse anti-Ctbp2; BD Transduction Labs, used at 1:200), myosin-VI (rabbit anti-myosin-VI; Proteus Biosciences, used at 1:200), anti-otoferlin (HCS-1, DSHB 1:500) and anti-GFP (NB100-1614, Novus Biologicals 1:500).

### Drug identification using LINCS and GDA

RNA-seq studies of cisplatin-resistant and -sensitive parental lines available in the public Gene Expression Omnibus (GEO) database were analyzed using the National Center for Biotechnology Information (NCBI)’s GEO2R tool (https://www.ncbi.nlm.nih.gov/geo/geo2r/) to identify differentially expressed genes between the two groups. Genes with an absolute log-fold change greater than 1 were downloaded from each study and analyzed with the GDA (http://gda.unimore.it/index.php) and LINCS (https://maayanlab.cloud/L1000CDS2/#/index) databases to identify compounds inducing similar gene expression profiles in various cell lines.

The LINCS analysis relies on a subset of the 1,319,138 genetic profiles originally compiled in the L1000 compendium (13). For each profile, an overlap score between 0-1 was given, indicating the fraction of genes overlapping from the gene set input. With over 500 identified compounds of interest, we further narrowed down the results of our screen by selecting those compounds with an overlap score >0.1, indicating at least a 10% overlap between the small molecule perturbation from the databases and our gene expression profile.

The GDA tool was also used to search the gene expression values of cells treated with 50,816 different compounds originally derived from the NCI-60 GI_50_ file (14). A P-value is generated for each identified drug based on the responsiveness of both parental and mutant cancer cell lines treated with each compound. The benefit of GDA lies in its comprehensive list of compounds in combination with over 73 cell lines. However, GDA requires separate inputs for up- and down-regulated genes, meaning that it does not provide profiles which comprehensively match differential gene expression. Compounds with a P-value <0.05 from GDA were selected from each database for further characterization.

### Signaling pathway analysis of cancer cell lines

The up and downregulated genes across all nine studies obtained by using the GEO2R tool were compiled, and duplicates were removed to exclude genes that had contradicting expression across the GEO studies. The resulting list of genes for all cancer lines was loaded into ShinyGO v0.66 (http://bioinformatics.sdstate.edu/go/) for pathway enrichment analyses. The ShinyGO analysis tool contains a total of 4,559 upregulated and 5,141 downregulated genes that can be used for gene set enrichment analysis (GSEA) to identify significantly enriched up and downregulated pathways. The gene expression data sets identified in our study were annotated using the KEGG database, and those with false detection rate (FDR) P-value <0.05 were reported.

### RNA-seq analysis of HEI-OC1 cells treated with cisplatin

HEI-OC1 cells (a generous gift from Dr. Kalinec, House Ear Institute) (18) were exposed to 70 µM cisplatin for 15 hours and total RNA was extracted with TRIzol (Thermo Fisher Scientific, USA). Samples (1 µg total RNA per sample, n=2 per treatment) were shipped to Novogene (California, USA) for RNA sequencing and bioinformatic analysis. The metadata file, raw sequencing fastq files and normalized expression values (in Excel format) are available from the NCBI GEO submission number: GSE180141.

### Cell culture and apoptosis assay

The apoptosis assay was performed as previously described (43). Briefly, HEI-OC1 cells were pretreated with each of the 30 drug candidates at concentrations ranging from 2 nM to 40 μM one hour before co-incubation with cisplatin, 50µM for an additional 19 hours. Caspase-Glo 3/7 assay (Promega, Madison, WI) (5) was run in triplicate and results normalized to cisplatin-only and medium-only controls. The percent of caspase activity was used to determine the relative protective effect of each compound, calculated using the following formula:

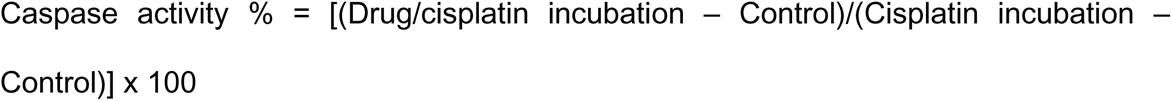

### Animals and drug administration

Animal procedures (fish and mice) were approved by the Institutional Animals Care and Use Committee at Creighton University.

For fish studies, 5 dpf larvae were maintained at 28.5°C in E3 water (5 mM NaCl, 0.17 mM KCl, 0.33 mM CaCl_2_, and 0.33 mM MgSO_4_, pH 7.2) and treated as previously described (54). HCs were counted from SO3 and O1-2 neuromasts.

For mouse studies, 5 to 7-weeks old FVB/NJ mice were obtained from The Jackson Laboratory (Bar Harbor, ME, USA), with a mix of males and females across experiments. FVB/NJ mice were treated with 10 mg/kg niclosamide via IP. Niclosamide was dissolved in 1% DMSO in normal saline (0.9% NaCl solution) and vortexed multiple times before injections. Niclosamide treatment started 24 hours before cisplatin (30 mg/kg divided into 2 doses, IP) or noise exposure (8-16 kHz octave band noise 100 dB SPL for 2 hours) and continued once daily for 3 more days. In the case of cisplatin, the administration protocol was designed based on a 5-days post-cisplatin treatment. Our previous publication showed that this *in vivo* cisplatin model produces similar results to the cisplatin models in which cisplatin was given in three cycles to CBA/CaJ mice (51).

### Zebrafish drug studies

For the screenings, 5-day post-fertilization (dpf) *Tg(brn3c:mGFP*) larvae were pre-incubated with each of the 30 drug candidates at 0.002, 0.0183, 0.165, 1.48, and 13.3 µM for as previously described (25, 54).

### Niclosamide/ezetimibe experiments in zebrafish

Synergistic interaction between niclosamide and the Nrf2 agonist ezetimibe was tested in 5-dpf zebrafish. Ezetimibe was used at 0.002 µM to 13.3 µM concentrations while niclosamide was used at 0.02 nM to 18.3 nM concentrations. Synergy analysis was conducted using Combenefit software that enables the analysis, advanced visualization and quantification of drug and other agent combinations. Combenefit performs combination analyses using the standard Loewe, Bliss, HSA and a newly developed SANE model (33, 34).

### Auditory brainstem response (ABR), distortion product otoacoustic emissions (DPOAEs) and noise exposure

FVB/NJ mice (6 to 7-weeks old) were used for the cisplatin and noise experiments and hearing function (ABRs and DPOAEs) tested as described before (5, 43). Briefly, ABR tests were performed 2-3 days before cisplatin exposure, and 5 days post-cisplatin exposure. For the noise experiments, auditory tests were performed 2-3 days prior to noise exposure, and 14 days post noise. Following the final auditory function measurements, mice were euthanized, and cochleae were collected for morphological assessment.

### Image analysis

After the different treatments the inner ear was microdissected and stained for CtBP2 (ribbon synapses marker) or myosin-VI (HC marker) as previously described (5, 43, 51). The organ of Corti was imaged using a confocal microscope (Zeiss LSM 700) with an oil immersion objective (40x, numerical aperture 1.3) and a digital zoom of 1X. The total number of OHCs was calculated by counting the number of cells in the three rows of OHCs within a 100-μm length of the cochlea. For IHC ribbon synapse quantification, 3D (x-y-z-axis) images were scanned with the 2X digital zoom. Each immunostained presynaptic CtBP2 puncta was counted as a ribbon synapse (26, 56). Synaptic ribbons of ten consecutive IHCs distributed within the mid-basal frequency region were counted. For neuromast imaging, samples were analyzed under a Zeiss LSM 710 confocal microscope with an oil immersion objective of 63X (numerical aperture 1.4) and 2x digital zoom.

### High-performance liquid chromatography (HPLC)

To assess whether niclosamide chemically interacts with cisplatin, we performed HPLC analysis. Stock solutions of cisplatin and niclosamide 1 mg/mL were prepared and mixed at a ratio of cisplatin:niclosamide 1:1 and 1:10 in the final injecting solution. Niclosamide- and cisplatin-only were also run.

### Experimental design and statistical analysis

#### Cell studies

For sample-size estimation we based on previous published data from our laboratory (5). All cell culture experiments were run in triplicate, and each considered a biological replicate for the statistical analysis. Outliers were considered to be those samples with percent caspase activity <-50%, which indicates total cell die-off likely due to a technical error. One-way ANOVA of drug-treated cells versus cisplatin-treated cells was used to calculate significance, followed by Dunnett’s multiple comparison test. Exact p values and 95% confident intervals (CIs) are included in the source data for **Figure 3A**. Results are presented as mean +/- SD.

Data from the bulk RNA-seq of cisplatin treated cells is available under GEO accession GSE180141.

#### Zebrafish studies

Five fish were employed per treatment and 2-3 neuromasts (SO3, O1 and O2) were inspected per fish. Each neuromast was considered a biological replicate. No explicit power analysis was performed to compute the sample size for the zebrafish experiments, sample size was based on previous studies from our laboratory (54). Only one experiment was performed for the initial screening of all the compounds and results were expressed as percentage of protection respect to controls. To further assess niclosamide’s effect, two independent biological experiments, were performed and results were expressed as number of HCs per neuromast. Statistical analysis: One-sample t-test run by the Combenefit software (33, 34) was used for the synergy studies. One-way ANOVA followed by Dunnett’s multiple comparison test was performed employing GraphPad for the rest of the studies. Exact p values and 95% CIs are included in the source data for Figure 3B. Fish samples were coded and HCs counts were assessed by an operator that was blinded to the group treatment. Results were presented as mean +/- SD.

#### Mice studies

Each animal was considered a biological replicate. For audiometric measurements, eight animals were used per treatment. For HC counting 4 organs of Corti from 4 different animals were inspected. For CtBP2 counting, 5 organs of Corti from 5 different animals were assessed. In both cases, each tissue sample was considered an independent biological replicate. The group size was calculated based on an effect size of 0.5 at alpha = 0.05 with an effective power of 0.868 (G*Power) (5, 43). Comparisons between the treatments for ABR (cisplatin exposure) were performed using two-way ANOVA followed by Tukey’s multiple comparison test and the Sidak’s multiple comparison test for ABRs and DPOAEs for the noise studies. One-way ANOVA followed by Dunnett’s multiple comparison test was used for CtBP2 puncta and OHCs analysis. ABR/DPOAE thresholds, HC count, and CtBP2 puncta counts were determined by an independent observer who was blinded to the group of mice.

#### For all the experimental data

GraphPad Prism v8/9 was used for statistical analysis. Unless specified, no outliers were identified employing the GraphPad tool function. When possible, equal number of males and females were employed for the experiments. Randomization was used for the zebrafish and mouse experiments. Statistical significance was set at p-value ≤0.05.

## ACKNOWLEDGEMENTS

We thank Emma Malloy and Tal Teitz for their help on cancer cell lines, Zhuo Li and Kan Lin for cell culture, Molly Kubesh for pathway analysis, and Mrs. Xianghong Liu for technical support. We also want to thank members of the Translational Hearing Center at Creighton University and Drs. Sung-Ho Huh and Wallace Thoreson at the University of Nebraska Medical Center for their comments. We thank Drs. Silvio Bicciato and Jimmy Caroli at the University of Modena, Italy for their initial assistance in the use of the GDA database. This work was supported in part by NIH-R01DC015444, NIH-R01DC015010, USAMRMC-RH170030, ONR-N00014-18-1-2507, and LB692/Creighton to JZ, by DoD-RH190050 and the Bellucci Foundation Award to MZ, by NIH-R43 DC018762 to PS (and subcontract to JZ), and by NIH1P20GM139762.

---

**Figure 1-table supplement 1.**
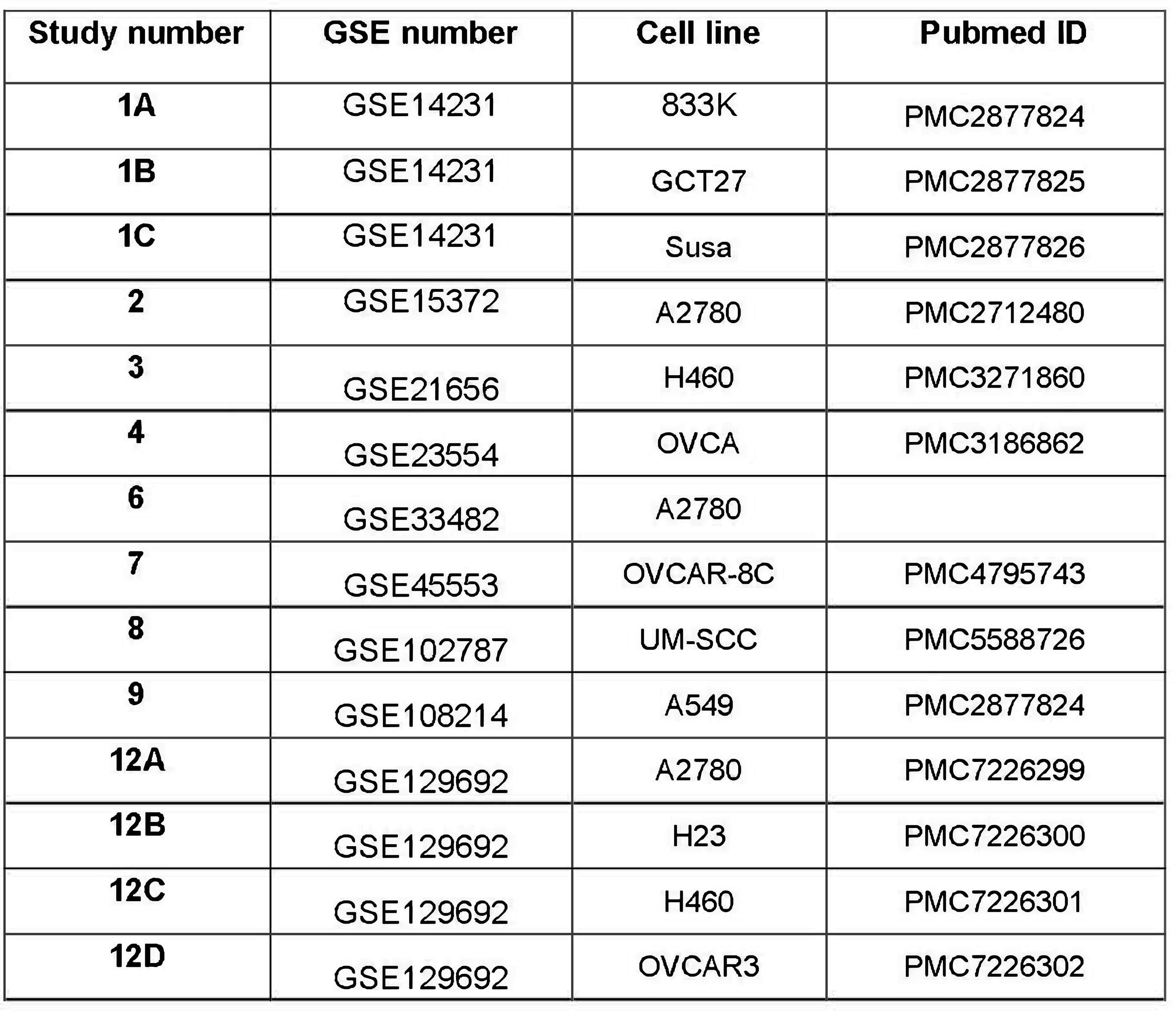
GEO studies of cisplatin-resistant cancer cell lines.

**Figure 2-figure supplement 1.**
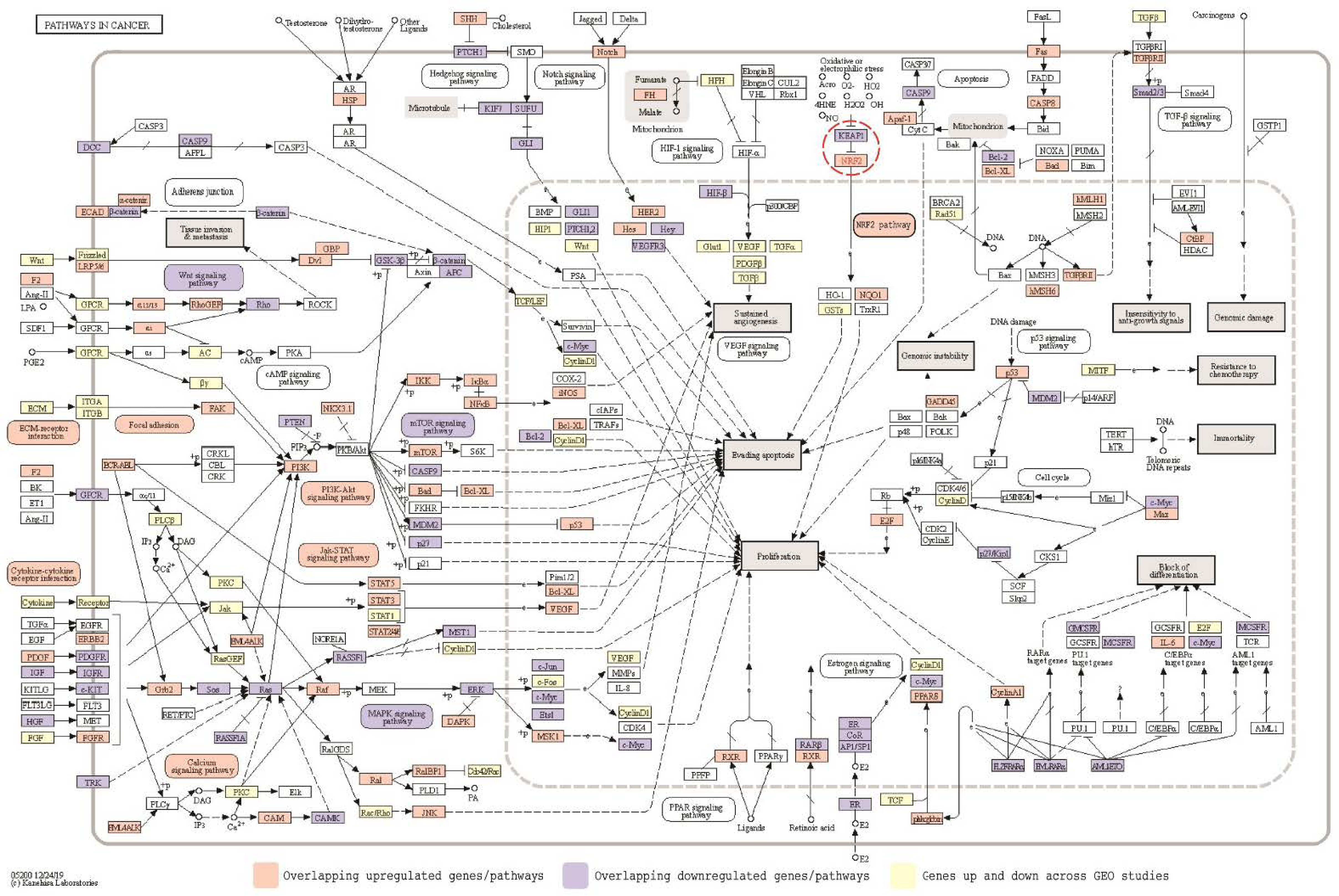
GSEA analysis of KEGG enriched molecular pathways of cisplatin-resistant cancer cells showed overlap with several pro-survival pathways. Overlapping DEG profiles in the cisplatin­resistant cancer cell lines and enrichment in pro-survival pathways, including the Nrf2 pathway (red circle), suggests several molecular mechanisms underlying cisplatin resistance.

**Figure 2-figure supplement 2.**
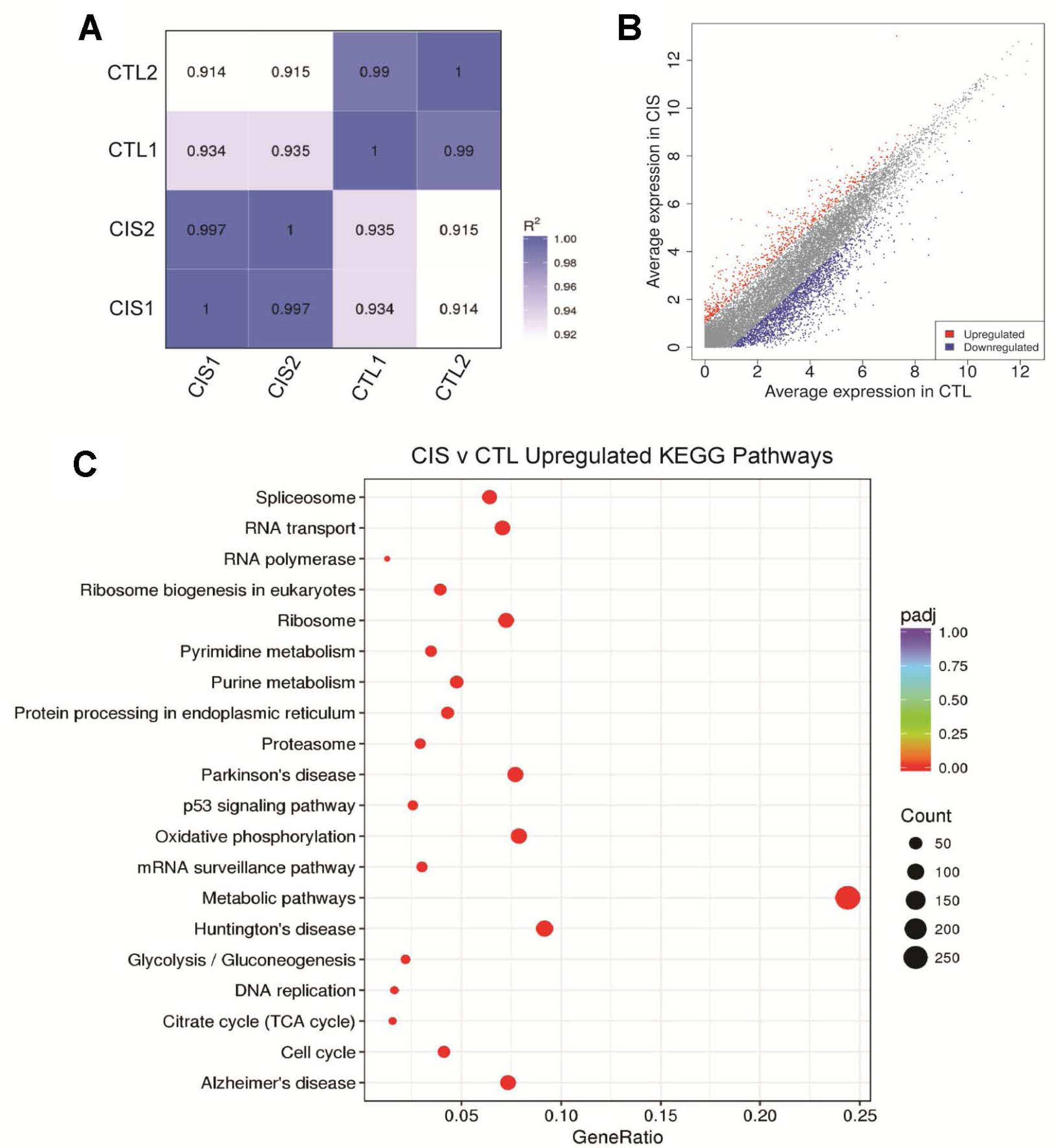
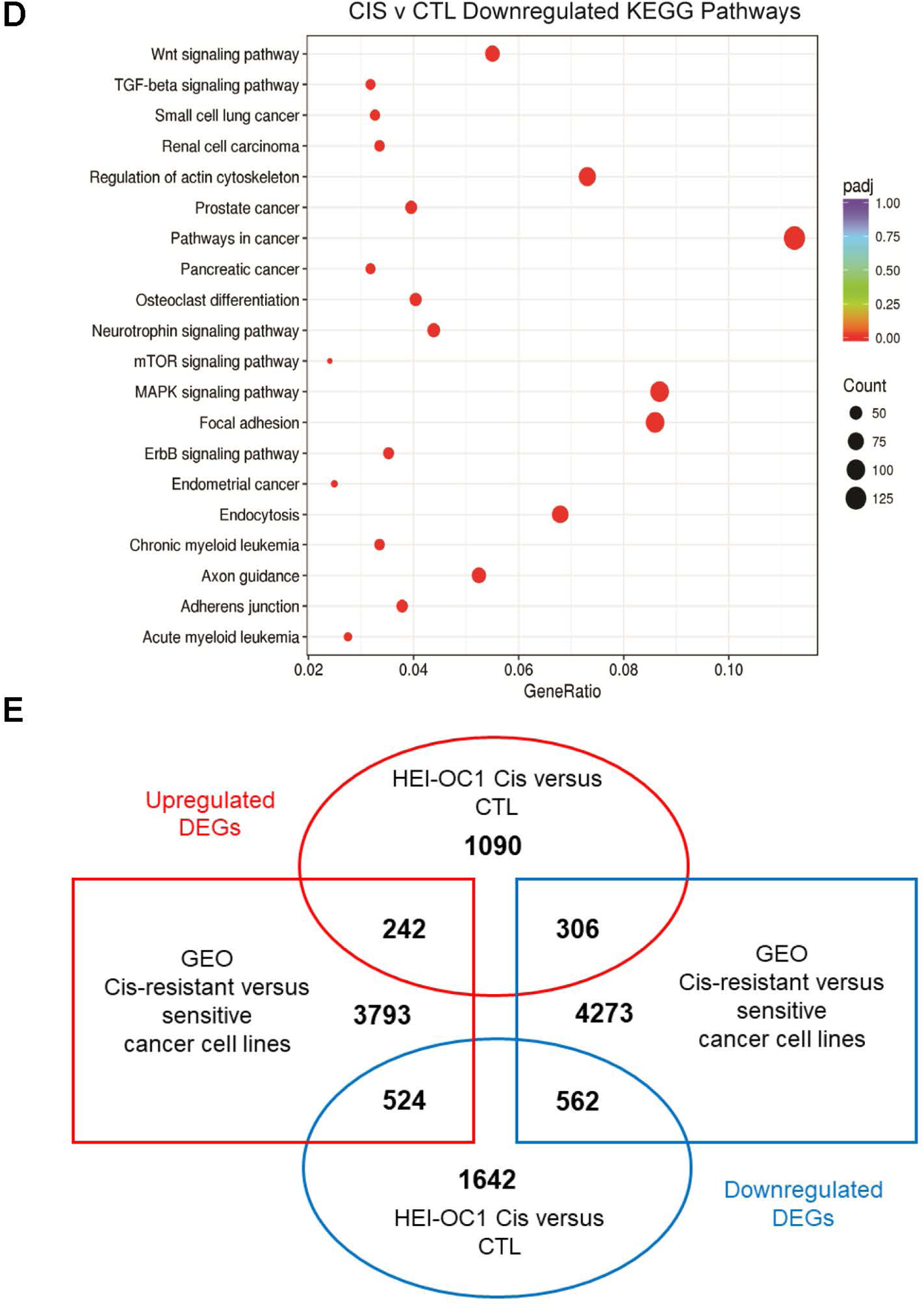
RNA-seq analysis of cisplatin-treated HEI-OC1 cells. (A) Pearson’s correlation matrix of cisplatin-treated (CIS) and untreated control (CTL) samples. (B) Scatter plot of differentially expressed genes in cisplatin-treated and control HEI-OC1 cells. (C) Upregulated and (D) downregulated KEGG pathways in cisplatin-treated HEI-OC1 cells. X-axis represents the ratio of differentially expressed genes to all themes annotated for this gene ontology (GO) term. Color scale represents the adjusted p-value (padj). Dot size represents the number of differentially expressed genes associated with that particular GO term. (E) Representation of up- and down-regulated gene overlaps from the GEO cancer cell data and HEI-OC1 cells.

**Figure 2-table supplement 2.**
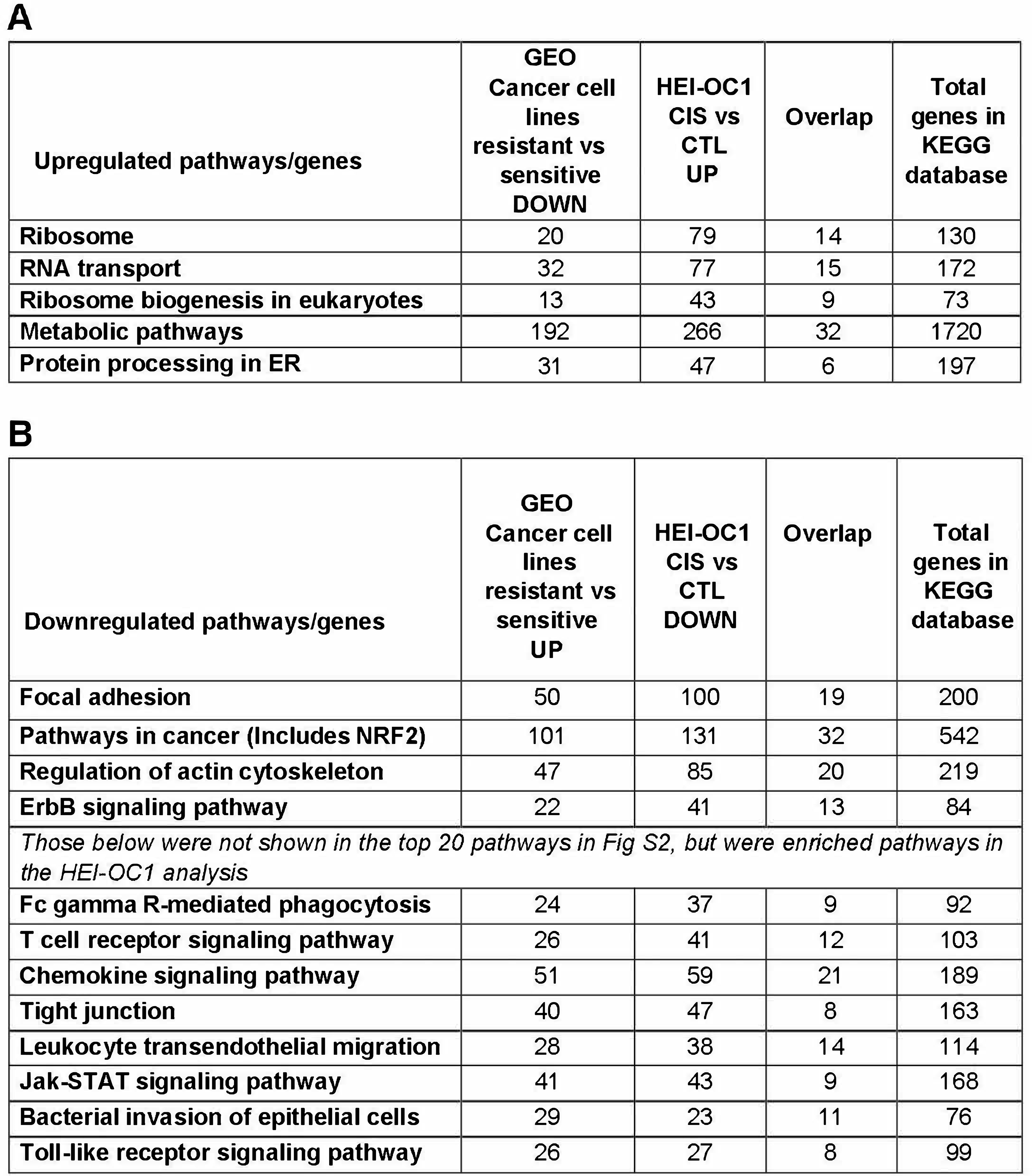
Pathways and gene overlaps between GEO cancer cell lines and HEI-OC1. A) Comparisons of the pathways/genes that are downregulated in the cisplatinresistance cancer cell lines while upregulated in the cisplatin-treated HEI-OC1 cells. B) Comparison of the pathways/genes that are upregulated in the cisplatin-resistant cells but downregulated in the cisplatin-treated HEI-OC1 cells. These pathways are likely to confer resistance to cisplatin-induced cell death.

**Figure 3-figure supplement 1.**
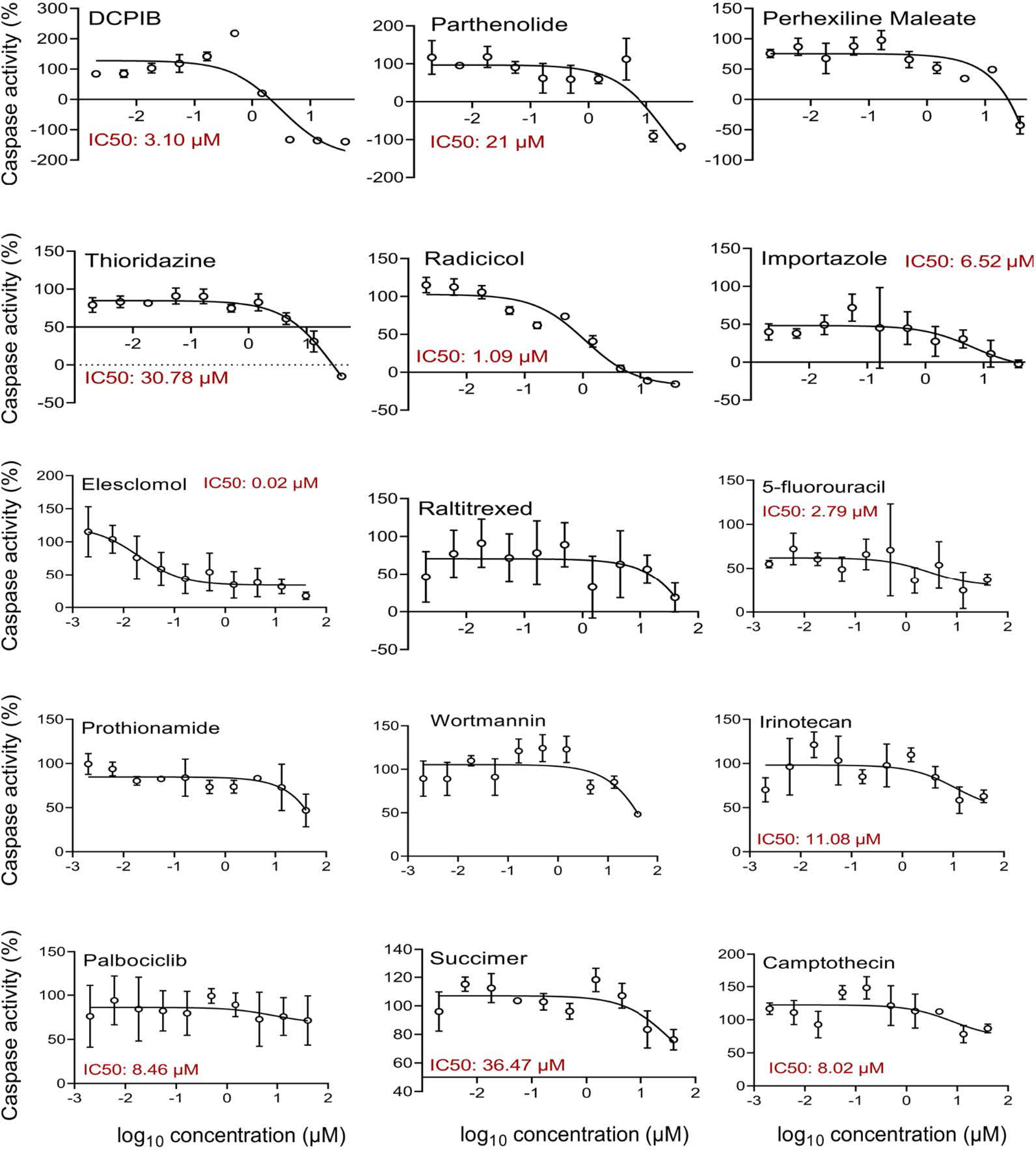
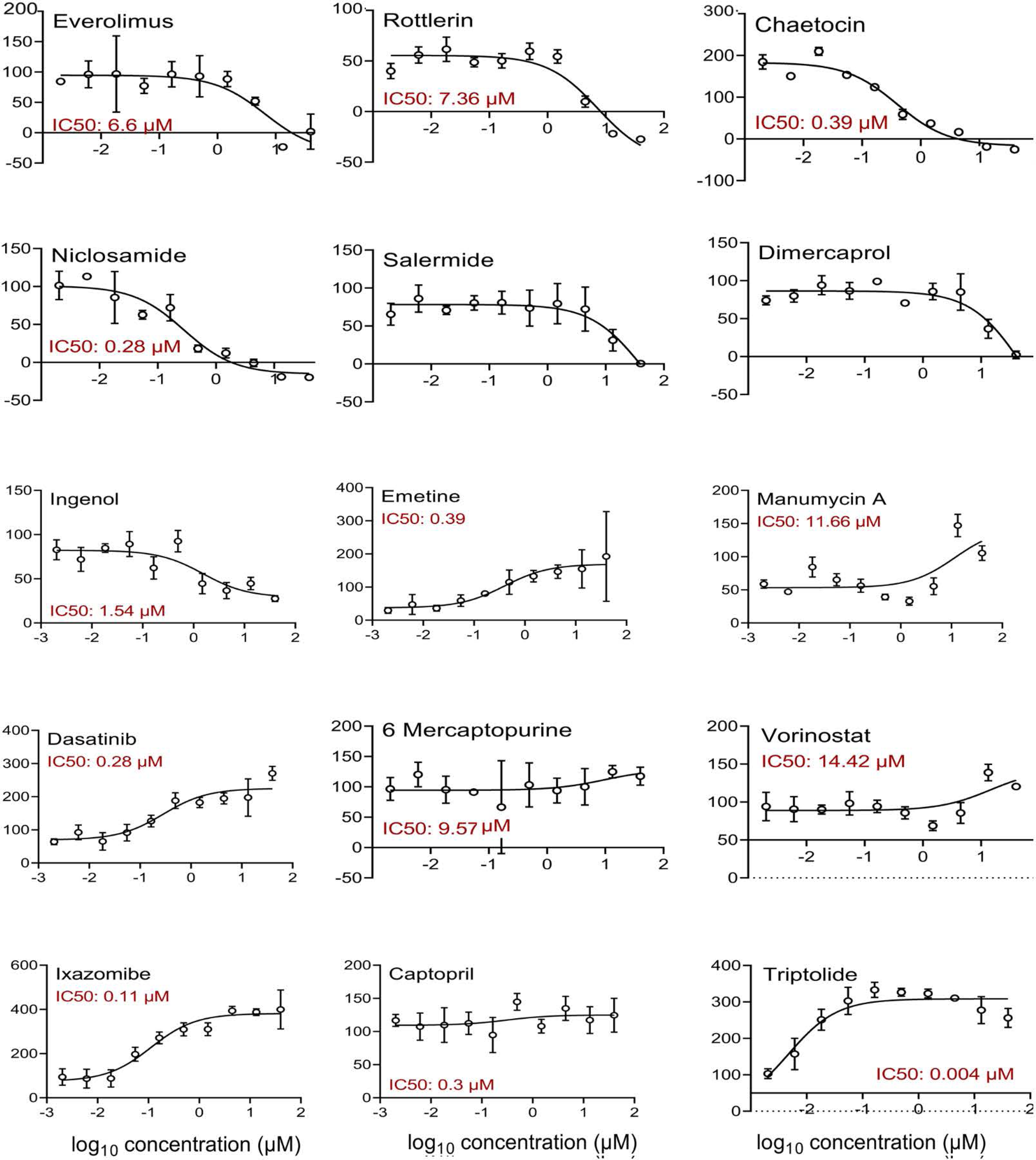
Dose-response curves for the top 30 experimental compounds. HEI-OC1 cells were exposed to cisplatin and various concentrations of the corresponding compounds. Caspase-3/7 activity was measured and plotted as a function of log10 compound concentration (µM). Caspase activity for all the treatments was normalized to cells treated only with cisplatin. Whenever possible, ICsos were calculated using GraphPad Prism software. Mean± standard error (n=3 per group).

**Figure 3-figure supplement 2.**
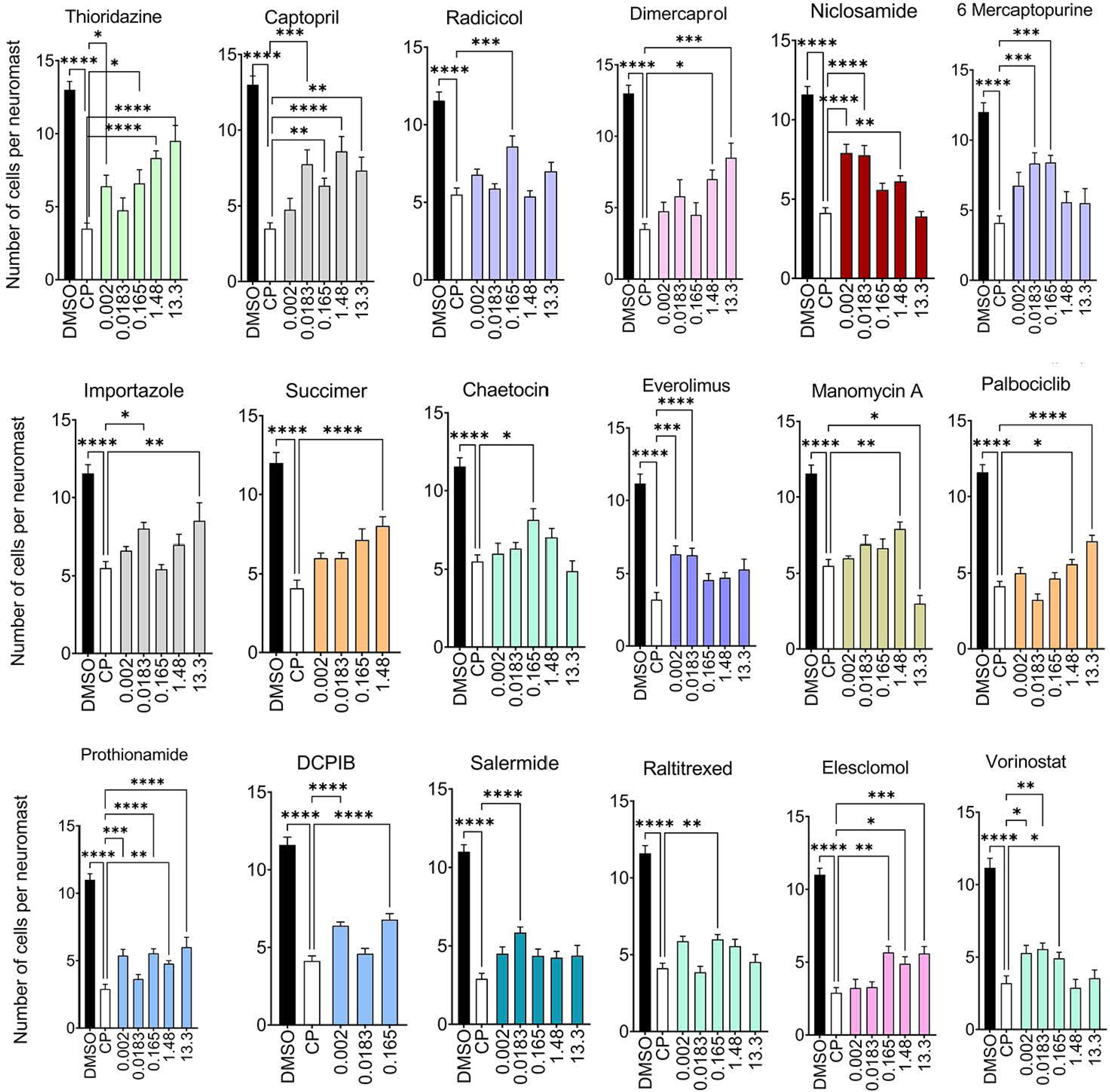
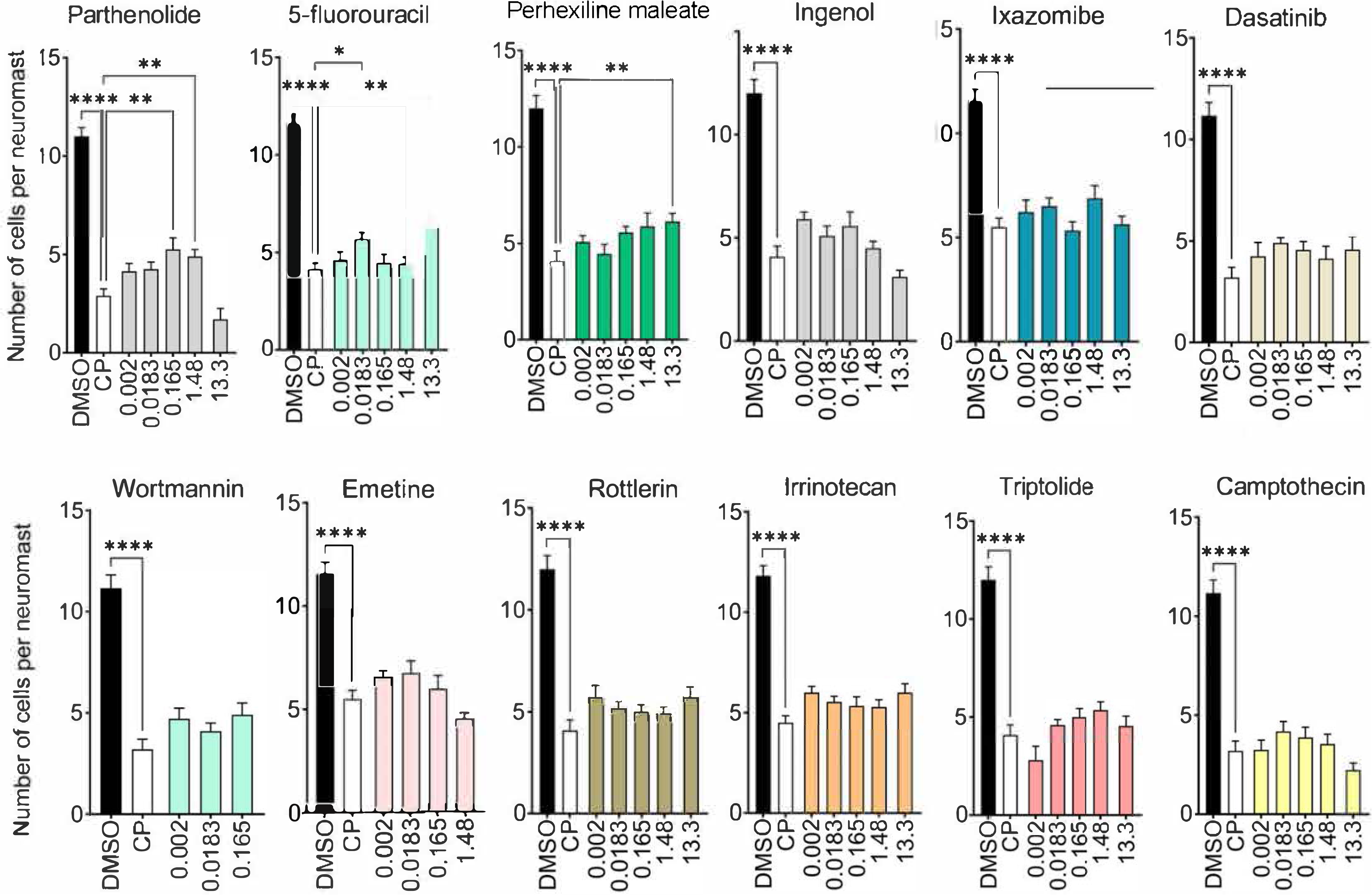
Characterization of the top 30 candidates in an *in vivo* model for cisplatin ototoxicity. Zebrafish were co-incubated with cisplatin and the 30 candidates at various concentrations as shown. Neuromast HCs were quantified and compared to cisplatin-only treated zebrafish. * P<0.05, **P<0.01, ***P<0.001, ****P<0.0001 versus cisplatin alone (one-way ANOVA followed by Dunnett’s multiple comparison test). Data shown as mean ± standard error (n^=^5 per group).

**Figure 3-figure supplement 3.**
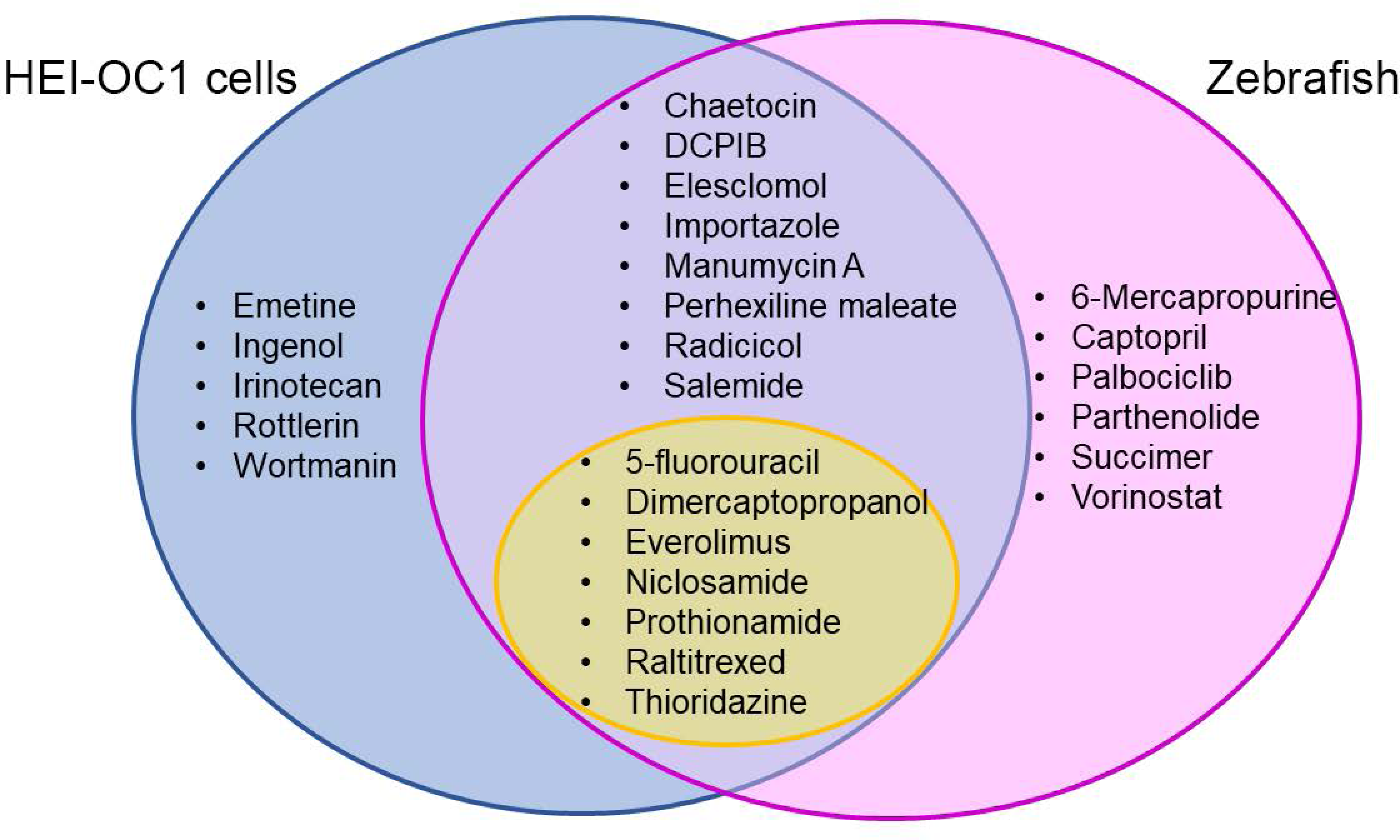
Analysis of the protective compounds identified in vitro in HEI-OC1 cells and in vivo in zebrafish experiments. Venn diagram showing the protective compounds identified in the two different screenings and the ones common to both assays. Experiments with HEI-OC1 identified 20 compounds with significant levels of protection (blue) while zebrafish experiments identified 21 compounds (pink). Fifteen compounds were commonly identified in both assays, with seven already approved by the FDA for other pathological conditions (yellow).

**Figure 3-figure supplement 4.**
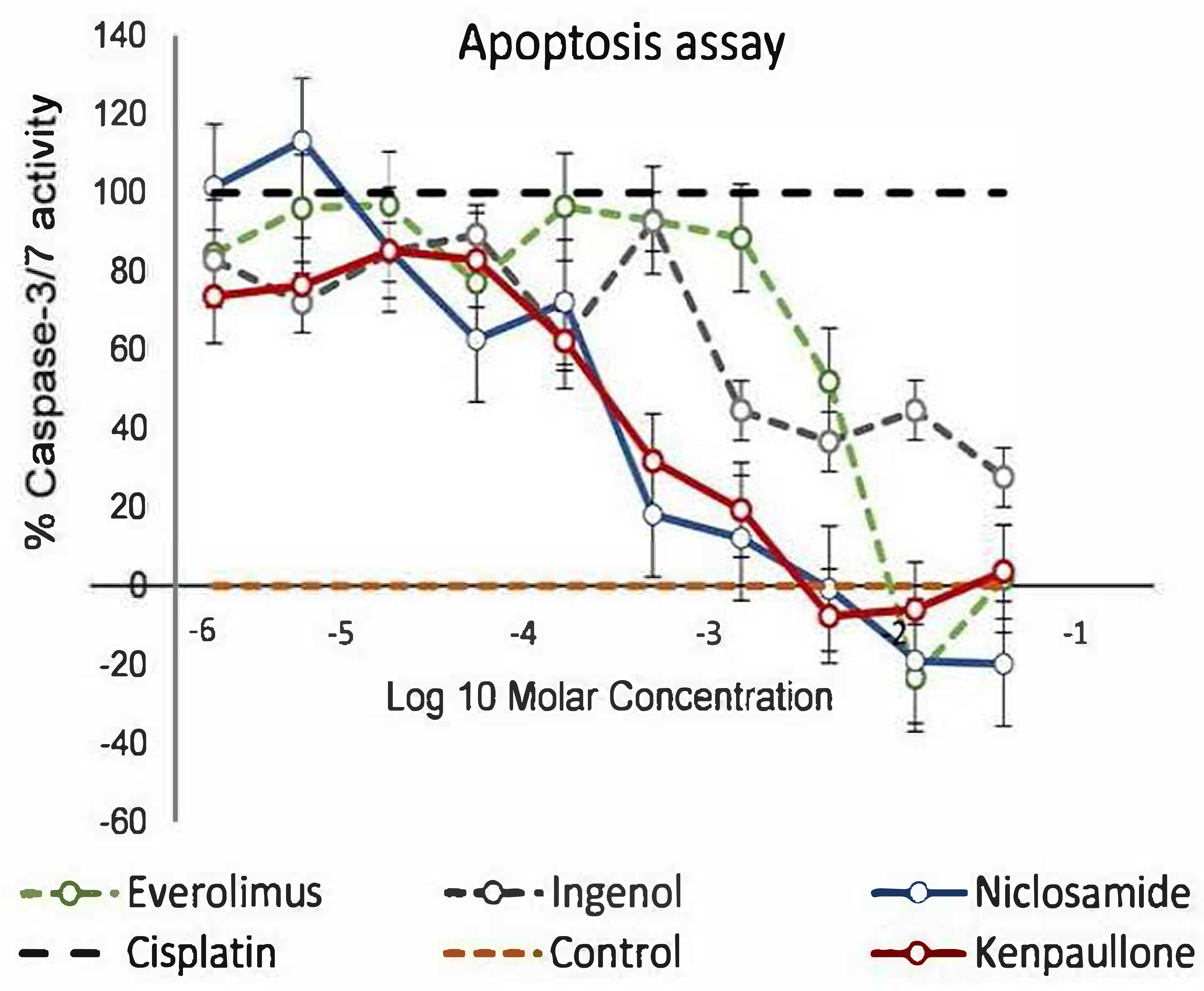
Comparison between niclosamide and everolimus, ingenol and kenpaullone. The protective effect of niclosam ide was com pared against two FDA-approved drugs ( everolimus and ingenol) as well as against kenpaullone that was identified in our previous screening.

**Figure 4-figure supplement 1.**
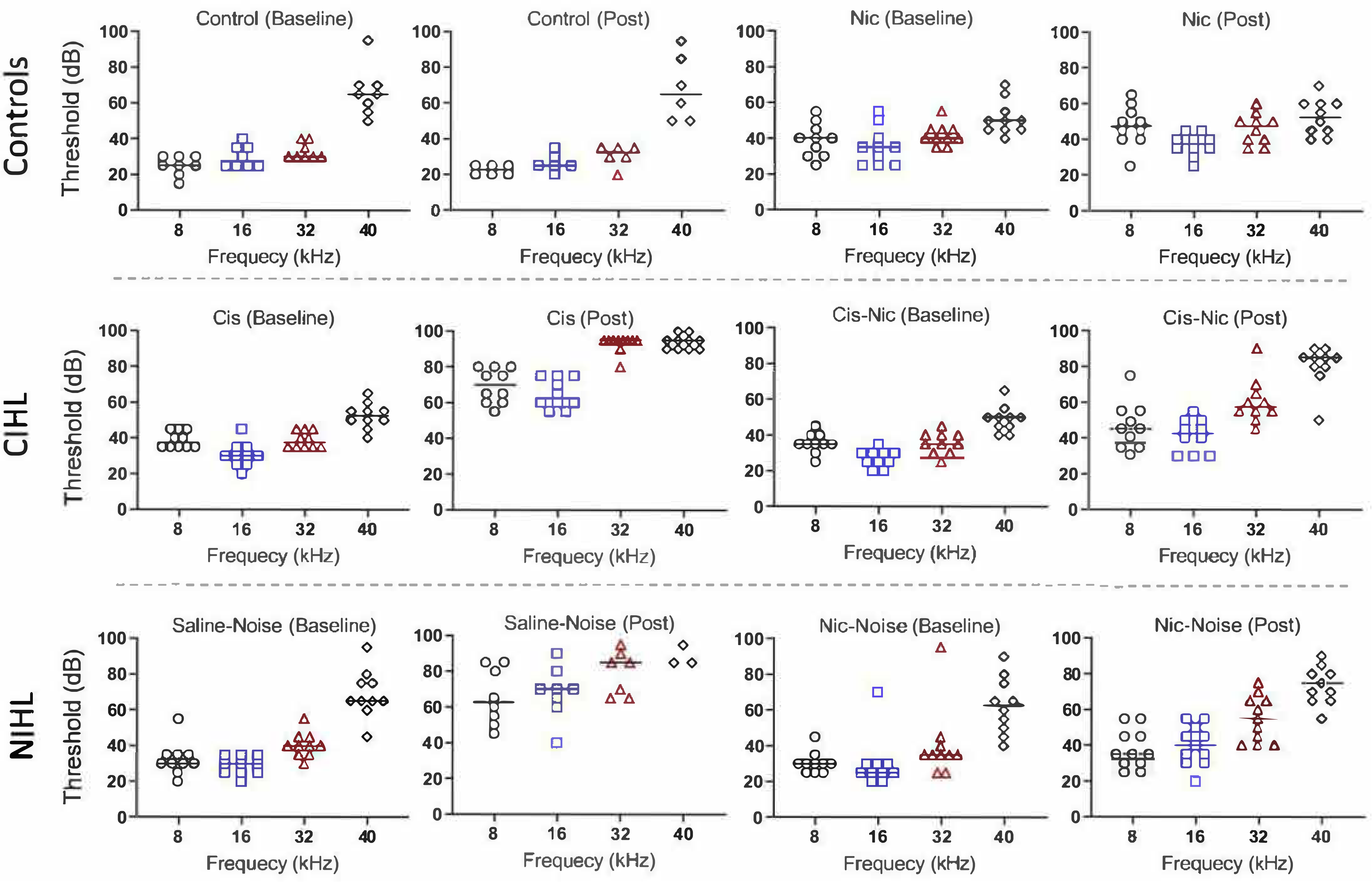
. Individual ABR thresholds. ABR thresholds from control (top row), cisplatin-treated (middle row), and noise-exposed (bottom row) animals at four different frequencies. Mice were also treated with vehicle or niclosamide 10 mg/kg for four consecutive days. Data shown as mean± standard error.

**Figure 4-supplement figure 2.**
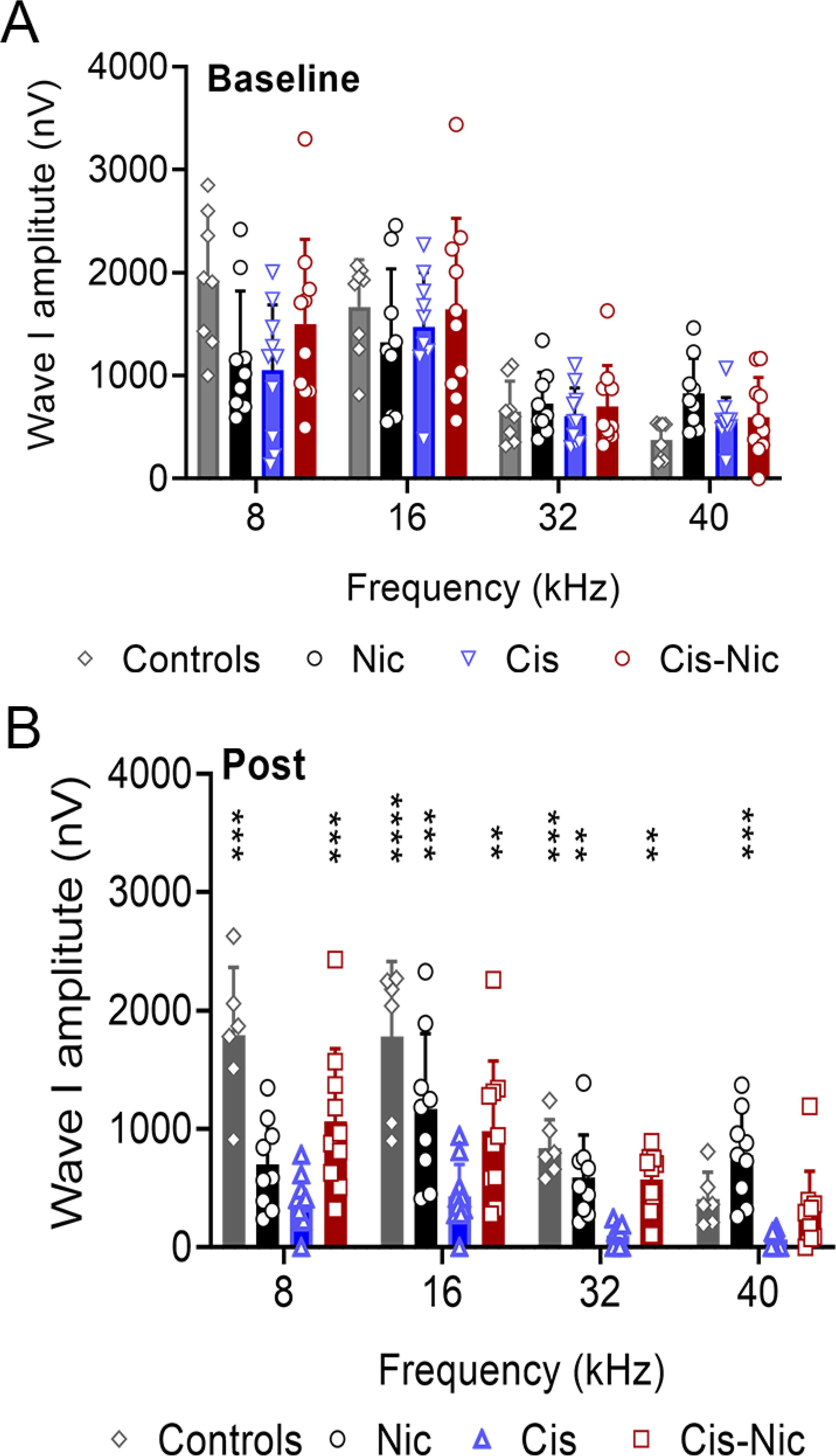
Niclosamide protects against cisplatin *in vivo*. Wave I amplitudes at baseline (A) showed no differences across all four groups. After cisplatin exposure (B), niclosamide was found to significantly increase wave I amplitudes from 8-32 kHz as compared to cisplatin-only treated mice (n=8 per group, **P<0.01, ***P<0.001, ****P<0.0001 versus cisplatin treatment, One Way ANOVA). Data shown as mean ± SD.

## REFERENCES

1. X. Chen, Y. Wu, H. Dong, C. Y. Zhang, Y. Zhang. Platinum-based agents for individualized cancer treatment. Mol. Med. 13, 1603–1612 (2013).

2. T. Magnes, A. Egle, R. Greil, T. Melchardt. Update on squamous cell carcinoma of the head and neck: *ASCO annual meeting*. 10, 220–223 (2017).

3. D. T. Dickey, Y. J. Wu, L. L. Muldoon, E. A. Neuwelt. Protection against cisplatin-induced toxicities by N-acetylcysteine and sodium thiosulfate as assessed at the molecular, cellular, and in vivo levels. J. Pharmacol. Exp. Ther. 314, 1052–1058 (2005).

4. H. S. So, C. Park, K. J. Kim, J. H. Lee, S. Y. Park, J. H. Lee, Z. W. Lee, H. M. Kim, F. Kalinec, D. J. Lim, R. Park. Protective effect of T-type calcium channel blocker flunarizine on cisplatin-induced death of auditory cells. Hear. Res. 204, 127–139 (2005).

5. T. Teitz, J. Fang, A. N. Goktug, J. D. Bonga, S. Diao, R. A. Hazlitt, L. Iconaru, M. Morfouace, D. Currier, Y. Zhou, R. A. Umans, M. R. Taylor, C. Cheng, J. Min, B. Freeman, J. Peng, M. R. Roussel, R. Kriwacki, R. K. Guy, T. Chen, J. Zuo. CDK2 inhibitors as candidate therapeutics for cisplatin- and noise-induced hearing loss. J. Exp. Med. 215, 1187–1203 (2018).

6. T. G. Baker, S. Roy, C. S. Brandon, I. K. Kramarenko, S. P. Francis, M. Taleb, K. M. Marshall, R. Schwendener, F. S. Lee, L. L. Cunningham. Heat shock protein-mediated protection against Cisplatin-induced hair cell death. J. Assoc. Res. Otolaryngol. 16, 67–80 (2015).

7. S. J. Kim, C. Park, A. L. Han, M. J. Youn, J. H. Lee, Y. Kim, E. S., Kim, H. J. Kim, J. K. Kim, H. K. Lee, S. Y. Chung, H. So, R. Park. Ebselen attenuates cisplatin-induced ROS generation through Nrf2 activation in auditory cells. Hear. Res. 251, 70–82 (2009).

8. Brock PR, Maibach R, Childs M, K. Rajput, D. Roebuck, M. J. Sullivan, V. Laithier, M. Ronghe, P. Dall’lgna, E. Hiyama, B. Brichard, J. Skeen, M. E. Mateos, M. Capra, A. A. Rangaswami, M. Ansari, C. Rechnitzer, G. J. Veal, A. Covezzoli, L. Brugieres, G. Perilongo, P. Czauderna, B. Morland, E. A. Neuwelt. Sodium Thiosulfate for Protection from Cisplatin-Induced Hearing Loss. N. Engl. J. Med. 378, 2376–2385 (2018).

9. M. Ryals, R. J. Morell, D. Martin, E. T. Boger, P. Wu, D. W. Raible, L. L. Cunningham. The Inner Ear Heat Shock Transcriptional Signature Identifies Compounds That Protect Against Aminoglycoside Ototoxicity. Front. Cell. Neurosci. 12, 445 (2018).

10. Kitcher SR, Kirkwood NK, Camci ED, et al. ORC-13661 protects sensory hair cells from aminoglycoside and cisplatin ototoxicity. JCI Insight 2019;4. PMID: 31391343.

11. Lamb J, Crawford ED, Peck D, et al. The Connectivity Map: using gene-expression signatures to connect small molecules, genes, and disease. Science 2006;313:1929–35. PMID: 17008526.

12. Musa A, Ghoraie LS, Zhang SD, et al. A review of connectivity map and computational approaches in pharmacogenomics. Brief Bioinform 2018;19:506–23. PMID: 28069634.

13. Duan Q, Reid SP, Clark NR, et al. L1000CDS(2): LINCS L1000 characteristic direction signatures search engine. NPJ Syst Biol Appl 2016;2. PMID: 28413689.

14. Caroli J, Sorrentino G, Forcato M, Del Sal G, Bicciato S. GDA, a web-based tool for Genomics and Drugs integrated analysis. Nucleic Acids Res 2018;46:W148–W56. PMID: 29800349.

15. Le BL, Andreoletti G, Oskotsky T, et al. Transcriptomics-based drug repositioning pipeline identifies therapeutic candidates for COVID-19. Res Sq 2021. PMID: 33821262.

16. Krishnamoorthy P, Raj AS, Roy S, Kumar NS, Kumar H. Comparative transcriptome analysis of SARS-CoV, MERS-CoV, and SARS-CoV-2 to identify potential pathways for drug repurposing. Comput Biol Med 2021;128:104123. PMID: 33260034.

17. Ge SX, Jung D, Yao R. ShinyGO: a graphical gene-set enrichment tool for animals and plants. Bioinformatics 2020;36:2628–9. PMID: 31882993.

18. Kalinec GM, Webster P, Lim DJ, Kalinec F. A cochlear cell line as an in vitro system for drug ototoxicity screening. Audiol Neurootol 2003;8:177–89. PMID: 12811000.

19. Ou HC, Santos F, Raible DW, Simon JA, Rubel EW. Drug screening for hearing loss: using the zebrafish lateral line to screen for drugs that prevent and cause hearing loss. Drug Discov Today 2010;15:265–71. PMID: 20096805.

20. Zhao J, He Q, Gong Z, Chen S, Cui L. Niclosamide suppresses renal cell carcinoma by inhibiting Wnt/beta-catenin and inducing mitochondrial dysfunctions. Springerplus 2016;5:1436. PMID: 27652012.

21. Tilabi J, Upadhyay RR. Adenoma formation by ingenol 3,5,20-triacetate. Cancer Lett 1983;18:317-20. PMID: 6406043.

22. Gregory MA, D’Alessandro A, Alvarez-Calderon F, et al. ATM/G6PD-driven redox metabolism promotes FLT3 inhibitor resistance in acute myeloid leukemia. Proc Natl Acad Sci U S A 2016;113:E6669–E78. PMID: 27791036

23. Le Prell CG, Yamashita D, Minami SB, Yamasoba T, Miller JM. Mechanisms of noise-induced hearing loss indicate multiple methods of prevention. Hear Res 2007;226:22–43. PMID: 17141991.

24. Sheth S, Mukherjea D, Rybak LP, Ramkumar V. Mechanisms of Cisplatin-Induced Ototoxicity and Otoprotection. Front Cell Neurosci 2017;11:338. PMID: 29163050.

25. Sheets L. Excessive activation of ionotropic glutamate receptors induces apoptotic hair cell death independent of afferent and efferent innervation. Sci Rep 2017;7:41102. PMID: 28112265.

26. Kujawa SG, Liberman MC. Acceleration of age-related hearing loss by early noise exposure: evidence of a misspent youth. J Neurosci 2006;26:2115–23. PMID: 16481444.

27. Chen W, Mook RA, Jr., Premont RT, Wang J. Niclosamide: Beyond an antihelminthic drug. Cell Signal 2018;41:89–96. PMID: 28389414.

28. Park JS, Lee YS, Lee DH, Bae SH. Repositioning of niclosamide ethanolamine (NEN), an anthelmintic drug, for the treatment of lipotoxicity. Free Radic Biol Med 2019;137:143–57. PMID: 31035006.

29. Sack U, Walther W, Scudiero D, et al. Novel effect of antihelminthic Niclosamide on S100A4-mediated metastatic progression in colon cancer. J Natl Cancer Inst 2011;103:101836. PMID: 21685359.

30. Ge SX, Jung D, Yao R. ShinyGO: a graphical gene-set enrichment tool for animals and plants. Bioinformatics 2020;36:2628–9. PMID: 31882993.

31. Zhang W, Xiong H, Pang J, et al. Nrf2 activation protects auditory hair cells from cisplatin induced ototoxicity independent on mitochondrial ROS production. Toxicol Lett 2020;331:1–10. PMID: 32428544.

32. Lee D, Hoon Han D, Nam K, Park J, Kim S, Lee M, Kim G, Min B, Cha B, Lee Y, Sung S, Jeong H, Ji H, Lee M, Lee J, Lee H, Chun Y, Kim J, Komatsu M, Lee Y, Bae S. Ezetimibe, an NPC1L1 inhibitor, is a potent Nrf2 activator that protects mice from diet-induced nonalcoholic steatohepatitis. Free Radic Biol Med 2016;99:520–32. PMID: 27634173

33. Loewe S. The problem of synergism and antagonism of combined drugs. Arzneimittelforschung 1953;3:285–90. PMID: 13081480.

34. Di Veroli GY, Fornari C, Wang D, et al. Combenefit: an interactive platform for the analysis and visualization of drug combinations. Bioinformatics 2016;32:2866–8. PMID: 27153664.

35. Organization DahlWH. https://www.who.int/news-room/fact-sheets/detail/deafness-andhearing-loss. 2020.

36. Le Prell CG. Otoprotectants: From Research to Clinical Application. Semin Hear 2019;40:162–76. PMID: 31036993.

37. Oh HC, Shim JK, Park J, et al. Combined effects of niclosamide and temozolomide against human glioblastoma tumorspheres. J Cancer Res Clin Oncol 2020;146:2817–28. PMID: 32712753.

38. Bhagat HA, Compton SA, Musso DL, et al. N-substituted phenylbenzamides of the niclosamide chemotype attenuate obesity related changes in high fat diet fed mice. PLoS One 2018;13:e0204605. PMID: 30359371.

39. Cerles O, Benoit E, Chereau C, et al. Niclosamide Inhibits Oxaliplatin Neurotoxicity while Improving Colorectal Cancer Therapeutic Response. Mol Cancer Ther 2017;16:300–11. PMID: 27980107.

40. Gratton MA, Eleftheriadou A, Garcia J, et al. Noise-induced changes in gene expression in the cochleae of mice differing in their susceptibility to noise damage. Hear Res. 2011; (1-2):211–26. PMID: 21187137

41. Jongkamonwiwat N, Ramirez MA, Edassery S, et al. Noise Exposures Causing Hearing Loss Generate Proteotoxic Stress and Activate the Proteostasis Network. Cell Rep. 2020; 24;33(8):108431. PMID: 33238128.

42. Lim N, Pavlidis P. Evaluation of connectivity map shows limited reproducibility in drug repositioning Sci Rep. 2021; 11(1):17624. PMID: 34475469

43. Ingersoll MA, Malloy EA, Caster LE, et al. BRAF inhibition protects against hearing loss in mice. Sci Adv 2020;6. PMID: 33268358.

44. Yu Y, Hu B, Bao J, et al. Otoprotective Effects of Stephania tetrandra S. Moore Herb Isolate against Acoustic Trauma. J Assoc Res Otolaryngol 2018;19:653–68. PMID: 30187298.

45. Salehi P, Akinpelu OV, Waissbluth S, et al. Attenuation of cisplatin ototoxicity by otoprotective effects of nanoencapsulated curcumin and dexamethasone in a guinea pig model. Otol Neurotol 2014;35:1131–9. PMID: 24841915.

46. A. A. Repurposing Existing Drugs for New Indications. The-Scientist 2017.

47. Malaviya AN. Landmark papers on the discovery of methotrexate for the treatment of rheumatoid arthritis and other systemic inflammatory rheumatic diseases: a fascinating story. Int J Rheum Dis 2016;19:844–51. PMID: 27293066.

48. Schweizer MT, Haugk K, McKiernan JS, et al. A phase I study of niclosamide in combination with enzalutamide in men with castration-resistant prostate cancer. PLoS One 2018;13:e0198389. PMID: 29856824

49. Burock S, Daum S, Keilholz U, Neumann K, Walther W, Stein U. Phase II trial to investigate the safety and efficacy of orally applied niclosamide in patients with metachronous or sychronous metastases of a colorectal cancer progressing after therapy: the NIKOLO trial. BMC Cancer 2018;18:297. PMID: 29544454.

50. Fernandez K, Spielbauer KK, Rusheen A, Wang L, Baker TG, Eyles S, Cunningham LL. Lovastatin protects against cisplatin-induced hearing loss in mice. Hear Res 2020;389:107905. PMID: 32062294.

51. Muniak MA, Rivas A, Montey KL, May BJ, Francis HW, Ryugo DK. 3D model of frequency representation in the cochlear nucleus of the CBA/J mouse. J Comp Neurol 2013;521:1510–32. PMID: 23047723.

52. Muniak MA, Rivas A, Montey KL, May BJ, Francis HW, Ryugo DK. 3D model of frequency representation in the cochlear nucleus of the CBA/J mouse. J Comp Neurol 2013;521:1510–32. PMID: 23047723.

53. Sergeyenko Y, Lall K, Liberman MC, Kujawa SG. Age-related cochlear synaptopathy: an early-onset contributor to auditory functional decline. J Neurosci 2013;33:13686–94. PMID: 23966690.

54. Zallocchi M, Hati S, Xu Z, Hausman W, Liu H, He D Z, Zuo J. Characterization of quinoxaline derivatives for protection against iatrogenically induced hearing loss. JCI Insights, 2021;6(5):e141561.

